# Zero-shot reconstruction of mutant spatial transcriptomes

**DOI:** 10.1101/2022.12.16.520397

**Authors:** Yasushi Okochi, Takaaki Matsui, Shunta Sakaguchi, Takefumi Kondo, Honda Naoki

**Author notes:** **Corresponding author** Graduate School of Integrated Sciences for Life, Hiroshima University, Kagamiyama 1-3-1, Higashihiroshima, Hiroshima 739-8526, Japan, **E-mail:**, **Co-corresponding author** Kyoto University Hospital, Shogoin Kawaracho 54, Sakyo, Kyoto, Kyoto 606-8397, Japan, **E-mail:**.

## Abstract

Mutant analysis is the core of biological/pathological research, and measuring spatial gene expression can facilitate the understanding of the disorganised tissue phenotype^1–5^. The large numbers of mutants are worth investigating; however, the high cost and technically challenging nature of experiments to measure spatial transcriptomes may act as bottlenecks^6^. Spatial transcriptomes have been computationally predicted from single-cell RNA sequencing data based on teaching data of spatial gene expression of certain genes^7^; nonetheless, this process remains challenging because teaching data for most mutants are unavailable. In various machine-learning tasks, zero-shot learning offers the potential to tackle general prediction problems without using teaching data^8^. Here, we provide the first zero-shot framework for predicting mutant spatial transcriptomes from mutant single-cell RNA sequencing data without using teaching data, such as a mutant spatial reference atlas. We validated the zero-shot framework by accurately predicting the spatial transcriptomes of Alzheimer’s model mice^3^ and mutant zebrafish embryos with lost Nodal signaling^9^. We propose a spatially informed screening approach based on zero-shot framework prediction that identified novel Nodal-downregulated genes in zebrafish. We expect that the zero-shot framework will provide novel phenotypic insights by leveraging the enormous mutant/disease single-cell RNA sequencing data collected.

## Introduction

Identifying spatial gene expression profiles is crucial for understanding whether a tissue of interest is functional in mutants and diseases. Recently developed spatially resolved transcriptomic technologies [*in situ* RNA capture for next-generation sequencing-based methods and *in situ* RNA sequencing for *in situ* hybridisation (ISH)-based methods]^1^ have enabled high-throughput measurement of gene expression profiles in a spatial context, providing valuable insights into the mechanism underlying tissue disorganisation in diseases^2–5^. However, many mutants are biologically and pathologically worth investigating. Nonetheless, the comprehensive measurement of the spatial transcriptomes of these mutants is limited by the cost and technically demanding nature of the technologies^10^. Moreover, these technologies often suffer from a trade-off between gene detection sensitivity and the number of genes measured^6^. In contrast, methods for computationally reconstructing spatial transcriptomes from single-cell RNA sequencing (scRNA-seq) data have a high gene detection sensitivity for whole transcriptomes^7^. According to the concept of reconstruction, dissociated scRNA-seq data are assembled by referring to teaching data of the spatial expression patterns of some genes, such as the ISH atlas. Many methods, including our previous method, Perler, have used various algorithms to reconstruct spatial transcriptomes from scRNA-seq data^11–19^. However, in mutant tissues, most of which have no spatial gene expression atlas, teaching data are generally unavailable, rendering the concept inapplicable.

Prediction without teaching data is a challenge for various tasks in image recognition and natural language processing that requires predicting previously unknown events. However, to compensate for the lack of teaching data, existing data, not including the teaching signals of interest, are trained as side information to refine the prediction. This concept, called ‘zero-shot learning’, has enormous potential for solving general prediction problems without teaching data, similar to how humans can predict a new event without ever experiencing it^8^.

In this study, we developed the first, to our knowledge, computational zero-shot framework (ZENomix) for the reconstruction of mutant spatial transcriptomes, without using teaching data, such as mutant spatial reference atlas. We leveraged the wild-type spatial reference atlas in our zero-shot system, easily accessible as side information. We reasoned that although the wild type had gene expression patterns different from those of mutant tissues, the underlying spatial coordinates of tissues were comparable when the wild-type and mutant tissue morphologies were similar. The wild-type reference atlas is used as a landmark point for spatial coordinates in tissues in ZENomix, helping in adding spatial information to mutant scRNA-seq data. ZENomix learns tissue spatial information by embedding a wild-type spatial reference atlas in the latent space and then mapping the mutant scRNA-seq data into the learned latent space. This latent spatial information is then used to reconstruct mutant spatial transcriptomes.

We first evaluated the performance of ZENomix in a mouse model of Alzheimer’s disease (AD) using simulated scRNA-seq data. We then used ZENomix to analyse mutant zebrafish early embryo scRNA-seq data. By comparing known ISH data for maternal-zygotic one-eyed pinhead (*MZoep*) mutants, we confirmed the spatial transcriptomes predicted by ZENomix. By predicting spatial gene expression using ZENomix, we identified previously unknown genes exhibiting spatially restricted gene expression changes, validated by conducting ISH experiments. These findings reveal that ZENomix provides a novel concept for zero-shot reconstruction of mutant spatial transcriptomes.

## Results

### Zero-shot reconstruction framework

ZENomix is a zero-shot learning computational method for predicting mutant spatial transcriptomes using wild-type spatial reference data as side information (**Fig. 1a**).

**Fig. 1.**
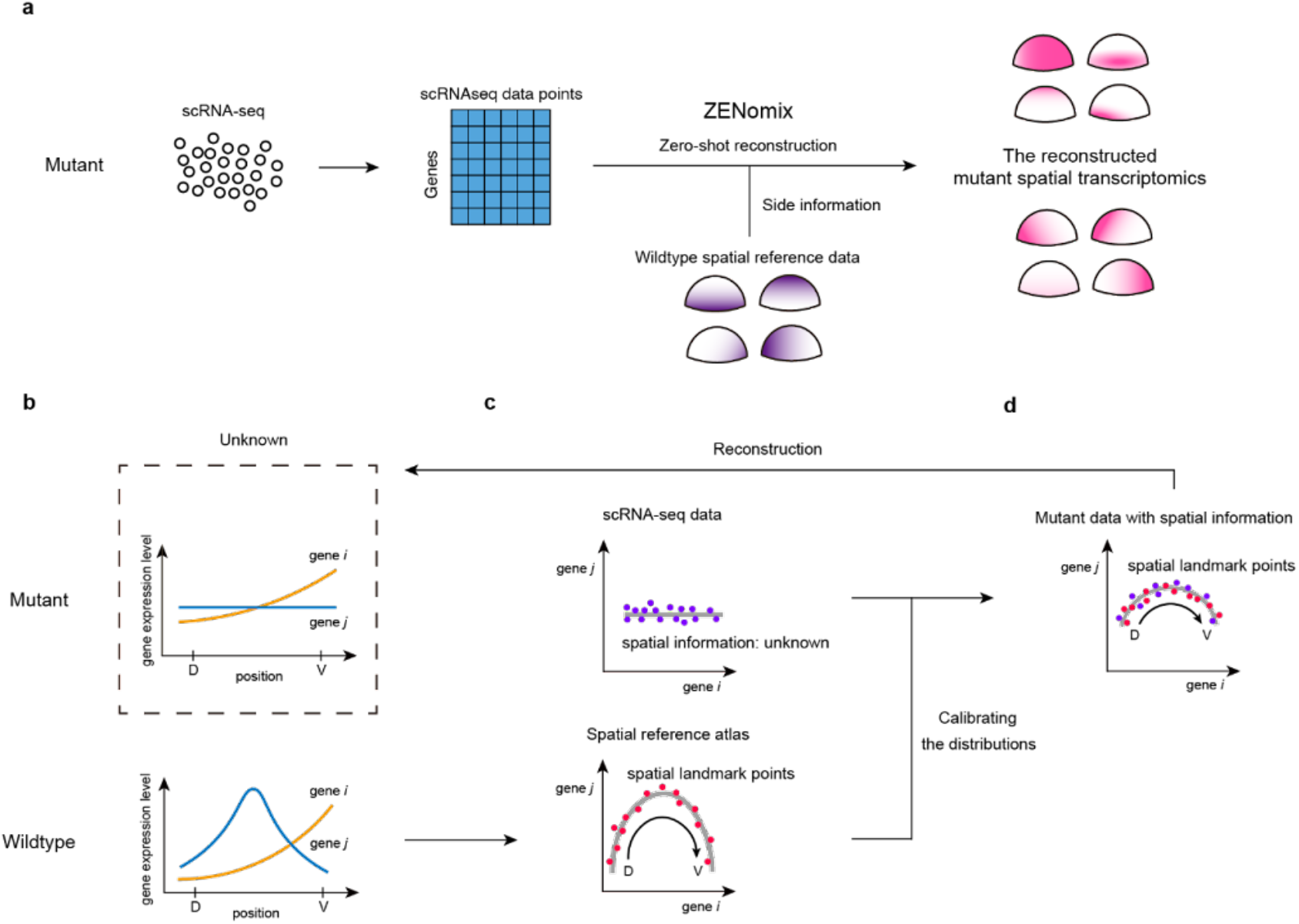
Zero-shot reconstruction concept. **a** Reconstruction of mutant spatial transcriptomes using ZENomix. **b**, Simple one-dimensional model of wild type and mutant tissues. The model tissues exhibited varying gene expression patterns along the dorsoventral (D-V) axis. Blue and orange lines indicate the expression profiles of genes i and j, respectively. **c**, Different trajectories in the gene expression space. Gene expression levels in mutants were measured using scRNA-seq, eliminating spatial information. In the wild type, gene expression levels were measured using *in situ* technology that has spatial information (black arrow) and are used as landmark points (postal codes) when adding spatial information to mutant scRNA-seq data. The red and blue points indicate the data points from the *in situ* technology and scRNA-seq, respectively. **d** Mutant gene expression data with spatial information. The trajectories were calibrated by matching the two data point distributions. After calibrating the differences in the trajectories, the lost spatial information of the mutant scRNA-seq data points can be retrieved by referencing landmark points having spatial information.

Predicting mutant spatial transcriptomes without using teaching data (i.e., a mutant spatial reference atlas) is generally challenging. To better understand the ZENomix framework, we started with a simple, one-dimensional tissue with two spatial gene expression profiles (e.g. dorsoventral axis patterning of the vertebrate neural tube by Shh and BMP/Wnt^20^) (**Fig. 1b**). The spatial information of each cell in this tissue could be expressed along its trajectory in the gene expression space (**Fig. 1c**). Furthermore, given that wild-type spatial reference data contain gene expression and spatial information, wild-type spatial reference data points can be used as landmarks for spatial information in gene expression spaces (**Fig. 1c**). This implies that if mutant scRNA-seq data points can be placed along this wild-type trajectory, their spatial information can be retrieved from landmark points possessing spatial information. However, in mutant tissues, the trajectory in the gene expression space was distorted because of varying gene expression profiles (**Fig. 1c**). Therefore, we calibrated the differences in these trajectories and retrieved the spatial information from the mutant scRNA-seq data by comparing cell distribution in the gene expression space (**Fig. 1d**).

Creating a zero-shot framework involves two steps: training and reconstruction. Using the abovementioned concept, the training step involves extracting the spatial information landmark points in the tissues from the wild-type spatial reference data. In practice, spatial reference data contain tens to thousands of gene expression profiles in two- or three-dimensional tissues; for example, 47 genes in zebrafish ISH data^11^ and 12,337 genes in mouse olfactory bulb (OB) spatial transcriptomics (ST) data^21^. To address the high-dimensional nature of these data, ZENomix embeds wild-type spatial reference data into the latent space to obtain spatial information landmark points using the Gaussian process latent variable model (GPLVM), a nonlinear dimensionality reduction method^22^ (**Fig. 2a**). The difference in distribution between the wild type and mutants was calibrated by minimising the distance between the two data distributions (**Fig. 2b**) using the maximum mean discrepancy (MMD) statistic^23^. The second step, reconstruction, was used to obtain the mutant spatial gene expression profiles. ZENomix mapped the landmark points derived from the wild-type data back to the mutant scRNA-seq space, and the mutant spatial transcriptomes were reconstructed as the weighted average of the mutant scRNA-seq data points using Gaussian process regression (arrows in **Fig. 2c**).

**Fig. 2.**
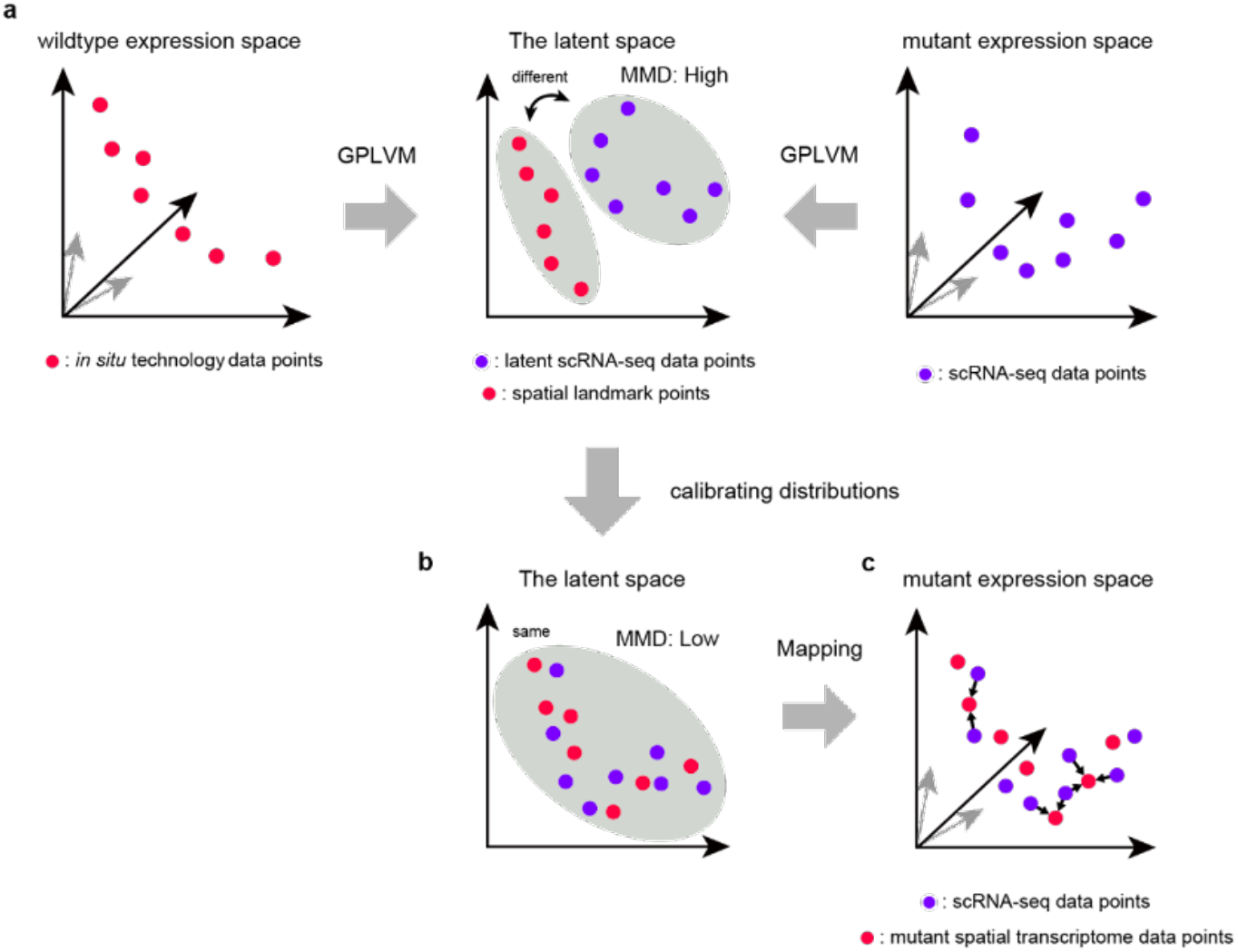
ZENomix data processing. **a**,**b**, First step of ZENomix. (**a**) Embedding into the latent space in ZENomix. In the wild-type expression space, there are high-dimensional *in situ* technology data points (red points; left panel); in the mutant expression space, there are high-dimensional scRNA-seq data points (blue points; right panel). ZENomix uses GPLVM to embed the wild-type data points into the latent space to obtain spatial landmark points. The mutant data points were also embedded in the latent space but with a data distribution different from that of the wild types. (**b**) Calibration of differences in the distributions. After mapping into the latent space, ZENomix calibrates the difference in the distributions of the two datasets by minimising the discrepancy between the distributions (MMD; see **Methods**). The grey-shaded region indicates the calibrated distribution. **c**, Second step of ZENomix. Following the first step, the spatial landmark points derived from wild-type data are mapped onto the mutant expression space by ZENomix. Mutant spatial transcriptomes (red points) were reconstructed using the weighted average of mutant scRNA-seq data points (blue points). Black arrows indicate the weights of the scRNA-seq data points.

To estimate the parameters of ZENomix, we proposed a new inference scheme (vGPLVM-MMD) by merging the variational GPLVM^24,25^ and MMD statistics (see **Methods**).

### ZENomix performance on simulated data

To determine whether our zero-shot reconstruction framework performed well, we used simulated scRNA-seq data from the triple transgenic AD (AD-mutant) mouse OB to evaluate ZENomix performance (**Fig. 3a**). These ST data represent gene expression in the mouse OB as obtained at points spatially arranged in a lattice (**Fig. 3b**). By concealing the actual spatial coordinates of the AD-mutant mouse OB data from Navarro et al., simulated scRNA-seq data (1,409 data points) were generated^3^. The ST data of the wild types from Ståhl et al.^21^ were used as spatial reference data (**Fig. 3c**). We used the ground-truth spatial coordinates of the simulated mutant scRNA-seq data as benchmarks.

**Fig. 3:**
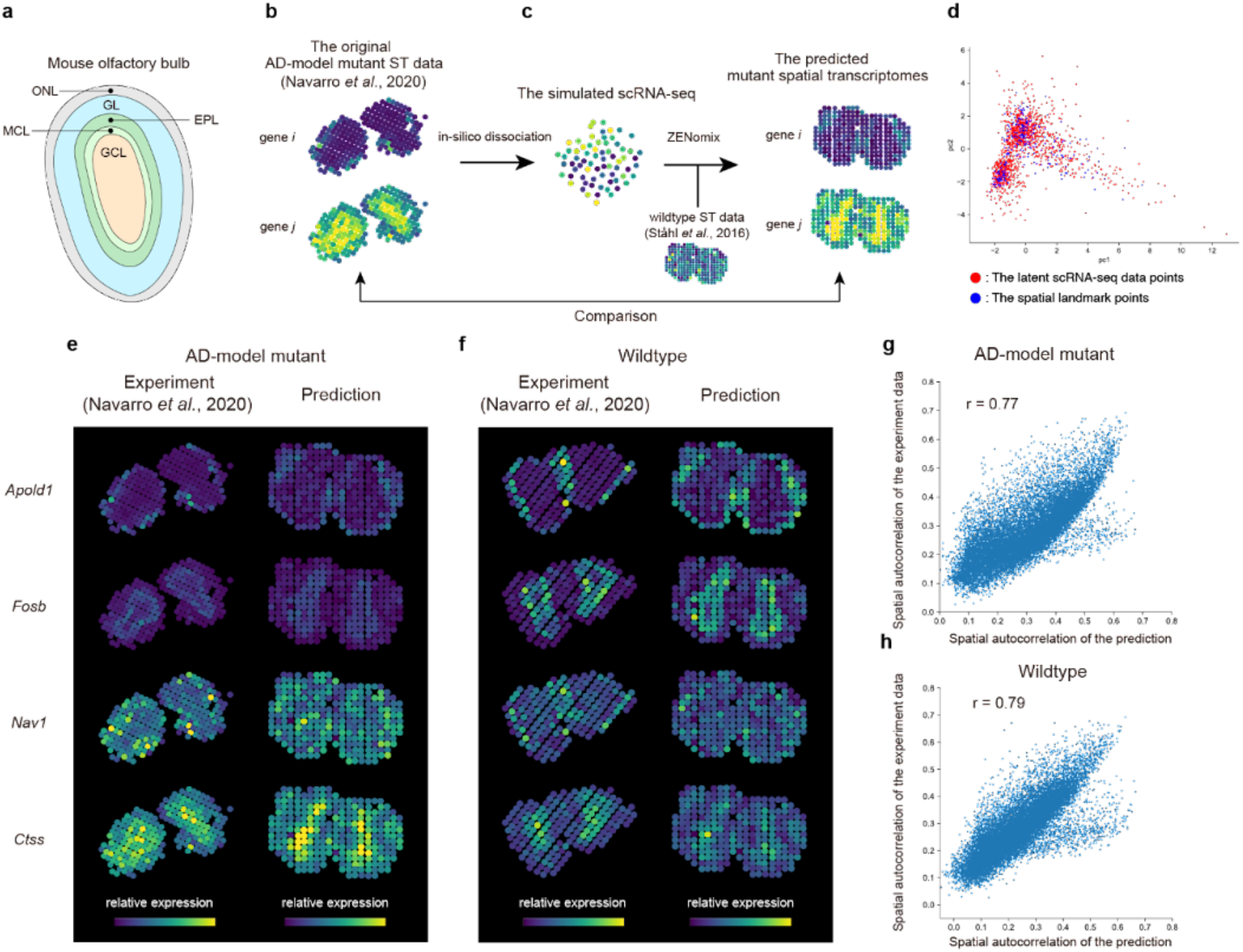
Zero-shot reconstruction of AD-mutant spatial transcriptomes. **a**, Anatomy of the mouse olfactory bulb^26^. GCL, MCL, EPL, GL, and ONL indicate granular, mitral, external plexiform, glomerular, and olfactory nerve layers, respectively. **b**,**c**, The experimental flow schematics. (**b**) Alzheimer’s disease mutant mouse olfactory ST data obtained from Navarro et al.^3^ *In silico* dissociation of the AD-mutant ST data yielded simulated scRNA-seq data. (**c**) Zero-shot reconstruction using ZENomix of AD-mutant spatial transcriptomes from Navarro et al. simulated AD-mutant scRNA-seq data.^3^ Wild-type ISH data from Ståhl et al.^21^ were used as side information. Prediction accuracy can be evaluated by comparing the predicted and original AD-mutant spatial transcriptomes. **d**, Scatter plot of the calibrated spatial landmark point and scRNA-seq data point distributions (Fig. 2b). Principal component analysis was used to visualise the latent space. **e**,**f** Original and predicted spatial transcriptomes of the (**e**) AD-mutant and (**f**) wild-type mouse olfactory bulb. **g**,**h** Comparison between the original and predicted spatial transcriptomes for the (**g**) AD-mutant and (**h**) wild-type mouse olfactory bulb. The original and predicted spatial profiles were evaluated quantitatively using Moran’s I value, a measure of spatial autocorrelation. Each dot indicates a gene (n = 16,037 and 16,046 genes for the AD-mutant and wild-type data, respectively).

We first confirmed that ZENomix could calibrate the distributions of two latent space data points (**Fig. 3d**) and that the model parameters converged (**Extended Data Fig. 1**). We showed that ZENomix successfully predicted the spatial transcripts of the AD-mutant mouse OB (**Fig. 3e**). We then performed the same analysis using simulated wild-type scRNA-seq data from the wild-type ST data of Navarro et al. (**Fig. 3f, Extended Data Fig. 2**). To assess the predictive accuracy of ZENomix for AD-mutant and wild-type mice, we compared the predicted spatial transcriptome with the original ST data by computing Moran’s I statistic, a measure of spatial autocorrelation, for all genes (n = 16,037 and 16,046 genes for AD-mutant and wild-type data, respectively), and found that the predicted and original spatial transcriptomes were correlated (r = 0.77 for the AD-mutant; r = 0.79 for the wild type; **Fig. 3g, h**). These findings validated the ability of ZENomix to execute zero-shot learning to reconstruct mutant spatial transcriptomes.

### Predicted spatial transcriptomes in zebrafish mutant

We used ZENomix to analyse the scRNA-seq data from mutant zebrafish embryos, for which the previous experimental method was challenging to apply because of the small, dome-shaped tissue. We used scRNA-seq data from an early embryo (at the 50% epiboly stage) of a maternal-zygotic one-eyed pinhead (*MZoep*)^27^ mutant obtained by Farrel et al. ^9^ The *MZoep* mutant lacks a Nodal signalling co-receptor, resulting in defects in the dorsal organiser, inducing mesoendodermal formation^28,29^ (**Fig. 4a**). For the spatial reference data, we used binary ISH data, including 47 genes obtained by Satija et al.^11^.

**Fig. 4:**
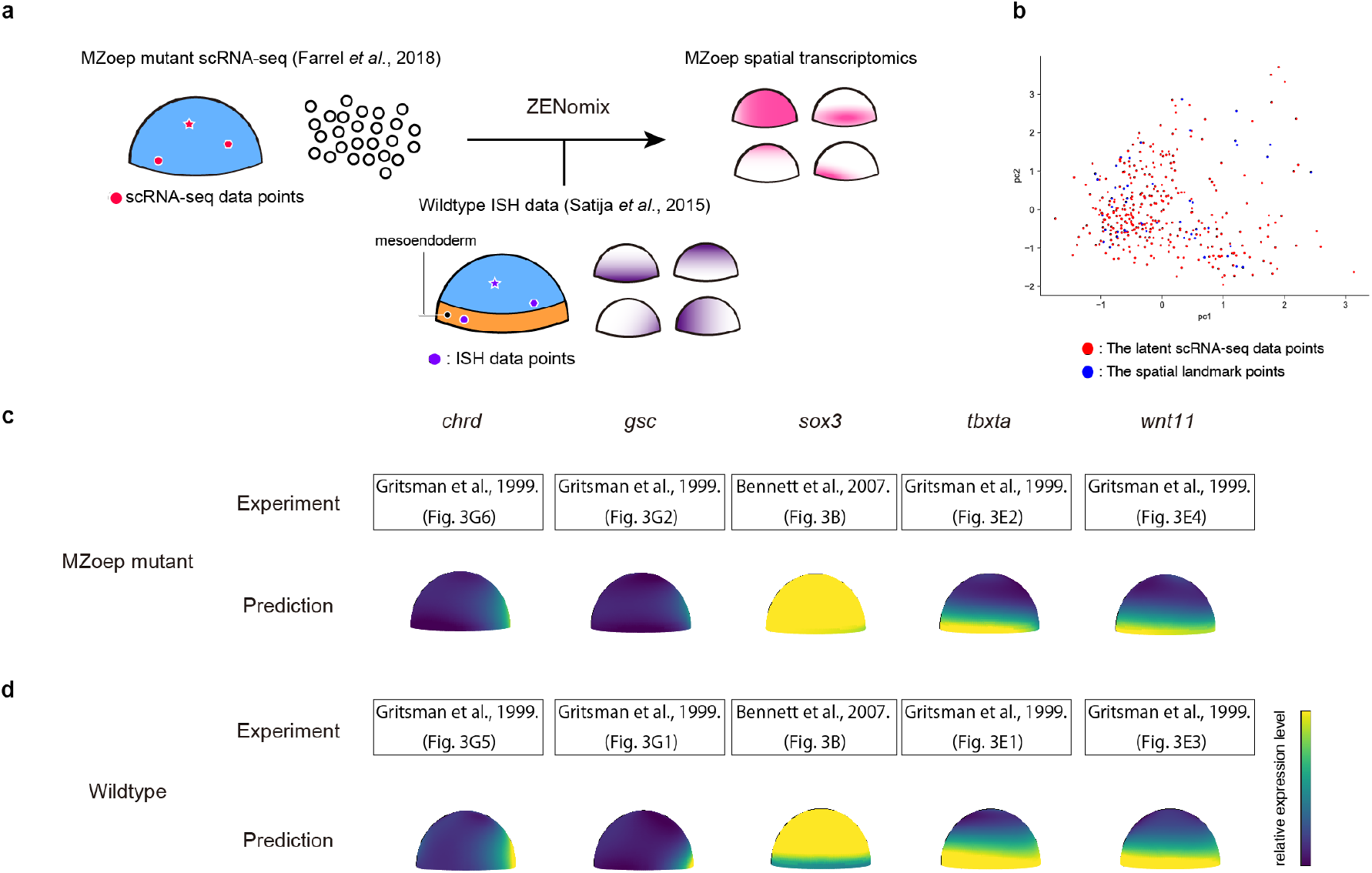
Spatial transcriptome prediction of zebrafish mutant embryos by ZENomix. **a**, Experimental flow schematic. ZENomix predicts the *MZoep*-mutant spatial transcriptomes from the mutant scRNA-seq data from Farrel et al.^9^ The wild-type ISH data from Satija et al.^11^ are used as side information. **b**, Scatter plot of the calibrated spatial landmark point and scRNA-seq data point distributions (corresponding to Fig. 2b). Principal component analysis was used to visualise the latent space. **c**,**d**, The experiment and prediction of the spatial transcriptomes of (**c**) the *MZoep*-mutant and (**d**) wild type. For the experiment, please see the ISH images of embryos at similar developmental stages reported by Gritsman et al.^27^ (*chrd, gsc, tbxta*, and *wnt11f2*; upper view) and Bennett et al.^30^ (*sox3*; lateral view).

First, we confirmed that ZENomix calibrated the distribution discrepancies between the two data points in latent space (**Fig. 4b, Extended Data Fig. 1–2**). The spatial transcriptomes of several genes known to be altered in *MZoep* mutants were predicted using ZENomix^27,30^, and all predictions were consistent with those in the previously published ISH images (**Fig. 4c-d**). Thus, ZENomix recapitulates the spatial gene expression changes caused by Nodal signalling mutations. These findings clearly indicated that ZENomix might successfully predict mutant spatial transcriptomes in a zero-shot manner.

### Identifying spatially differentially expressed (DE) genes

To identify the spatially DE genes in *MZoep* embryos, we compared the reconstructed mutant spatial transcriptome data with those of the wild type (**Fig. 5**). First, we computed the difference between the reconstructed spatial transcriptomes in the wild type and mutant (**Fig. 5a**). We then plotted the maximum absolute value and standard deviation of the expression changes in the *MZoep*-mutant transcriptome for each gene (n = 26,545 genes) and screened 142 Nodal-associated genes (**Fig. 5b**). We focused on expression changes for further screening in the embryo margin (red box in **Fig. 5c**), mostly affected by Nodal signalling defects. By plotting the average expression changes only in the embryo margin, the Nodal-associated genes were classified into two groups: 101 and 41 putative Nodal-upregulated (NU) and putative Nodal-downregulated (ND) genes, respectively (**Fig. 5c, Extended Data Fig. 3–5**).

**Fig. 5:**
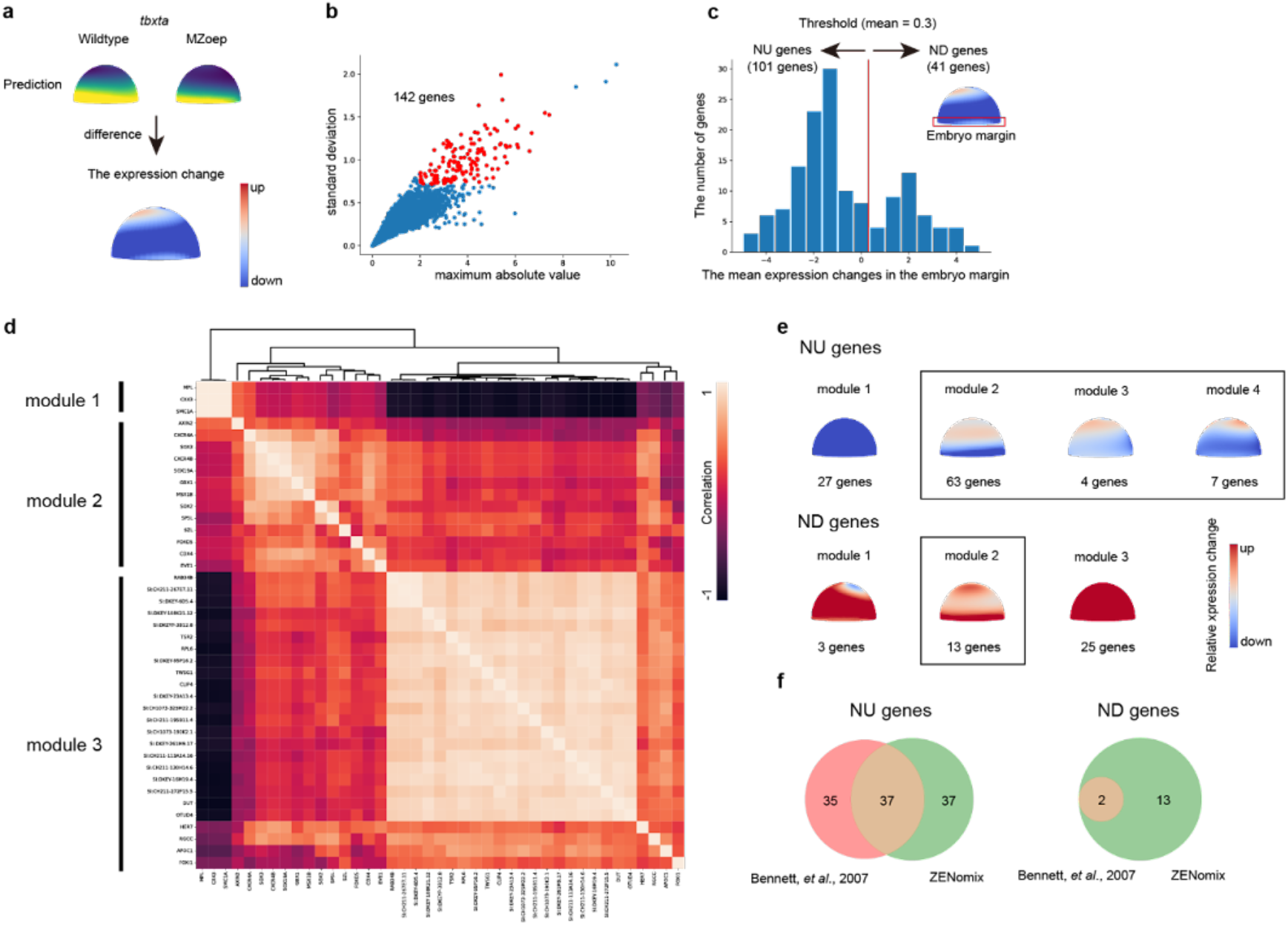
Screening of the putative spatially differentially expressed genes. **a**, Schematic depicting the expression changes calculated by subtracting the predicted spatial transcriptome of the wild type from that of the *MZoep*-mutant embryos. **b**, Gene screening scatter plot. The x- and y-axes indicate the maximum absolute value and standard deviation of the expression changes, respectively. Each dot indicates a gene (n = 26,545 genes). The red dots indicate selected genes (nodal-associated genes; n = 152). **c** Histogram of mean expression changes within the embryo margin. The red line (mean = 0.3) indicates the classification threshold between NU and ND genes. The red box in the inset indicates the embryo margin, within which the mean values were calculated. **d** Hierarchical clustering of putative ND genes. The heatmap indicates the correlation between gene expression changes among newly screened ND genes. **e**, The Average spatial gene expression changes in each module. Four and three modules for NU and ND genes are shown. The black rectangles indicate the modules included in further analysis. **f** Venn diagrams for sets of putative NU and ND genes screened by ZENomix and from a previous study by Bennett and colleagues^30^.

We then performed hierarchical clustering for each putative NU and ND gene to better understand Nodal-associated gene expression alterations (**Fig. 5d**). We identified four and three modules in putative NU and ND genes, respectively (**Fig. 5e, Supplementary Table 1**). Finally, we excluded the genes in module 1 of the putative NU genes and modules 1 and 3 of the putative ND genes since they showed universal gene expression changes in the whole embryo, yielding 87 putative spatially DE genes (74 putative NU genes and 13 putative ND genes). Notably, 50.0% (37/74) of these putative NU genes and 15.3% (2/13) of these putative ND genes were shared with 72 and two genes, respectively, identified in a similar bulk microarray screening in *MZoep* embryos^30^, suggesting that our screening was consistent with that of a previous study (**Fig. 5f**).

### New genes repressed by Nodal signalling

Nodal signalling is critical to induce and maintain the dorsal organiser and repress ectodermal cell fates^30^. Bennett et al. suggested that Nodal represses some target genes; nonetheless, only two genes (*sox2* and *sox3*) have been identified as downregulated via Nodal signalling^30^. Notably, ZENomix screening revealed 13 putative ND genes, including *sox2, sox3*, and 11 unknown genes (**Fig. 5f**).

To validate the candidate ND genes, we used ISH to determine whether the reconstructed spatial transcriptomes of the 11 unknown genes in *MZoep*-mutant embryos followed the spatial gene expression profiles by the ISH experiments. Since *MZoep* is not kept in our zebrafish facility, and *lefty1* overexpression, which encodes the Nodal inhibitor, can phenocopy *MZoep* mutants and *cyclops/squint* double mutants^31,32^, we used *lefty1*-overexpressed embryos instead of *MZoep* mutants for ISH evaluation of ZENomix predictions (*lefty1*-overexpressed embryos; see **Methods**). We found that among the 11 candidate genes, the predicted spatial gene expression patterns of eight genes (*cdx4, cxcr4a, cxcr4b, eve1, foxd5, sox19a, sp5l*, and *szl*) correlated with the ISH results, indicating that ZENomix discovered eight new genes repressed via Nodal signalling (**Fig. 6**).

**Fig. 6:**
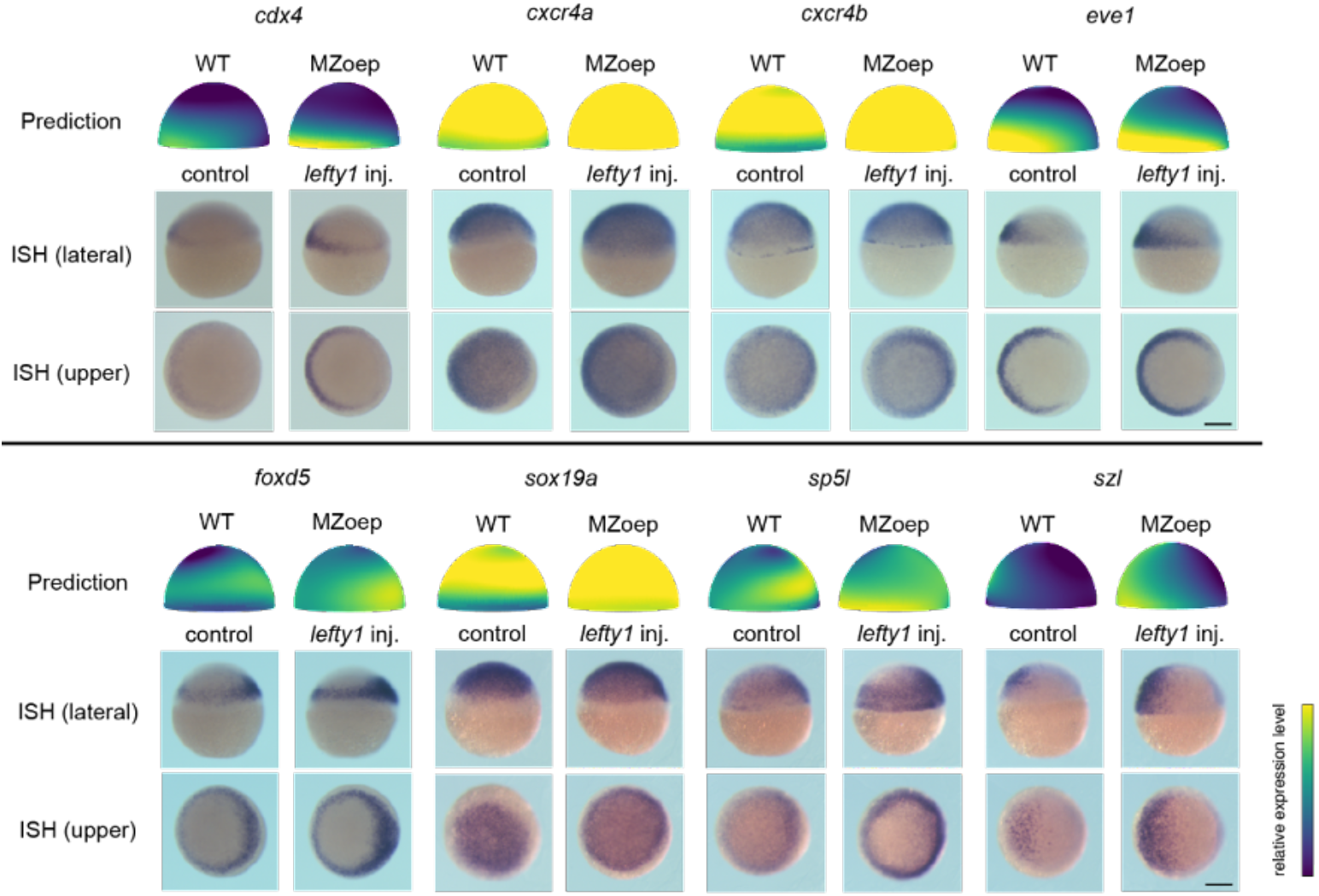
ISH validation of the Nodal-downregulated genes discovered from ZENomix-predicted mutant spatial transcriptomes. Whole-mount ISH experiments on eight identified ND genes. Top to bottom: ZENomix-predicted expression pattern, lateral ISH view, and upper ISH view. For each gene, wild type and mutant expression patterns are displayed (ZENomix prediction: wild type and *MZoep*-mutant embryos; ISH experiment: control and *lefty1*-injected embryos). Scale bar, 200 μm.

The expression patterns of the other three candidate genes (*axin2, gbx1*, and *msx1b*) were inconsistent with those of the ISH results (**Extended Data Fig. 6**). In the ISH assays, *axin2* specifically showed comparable ubiquitous expression patterns between wild type and *lefty1*-injected embryos, but the number of *axin2*-positive cells increased in *MZoep*-mutant embryos than in wild-type embryos in the scRNA-seq data (**Extended Data Fig. 6**). In contrast, *gbx1* and *msx1b* were unexpressed in either embryo in the ISH experiments, despite being expressed in the scRNA-seq data **(Extended Data Fig. 6)**. These inconsistencies may be partly attributed to expression property differences, such as measurement differences, batch effects, effects of mRNA injection, and phenotypic differences between *MZoep* and *lefty1* overexpression between *MZoep*-mutant and *lefty1*-overexpressed embryos. In addition, because *MZoep*-mutant embryos lack endoderm-derived tissue at a later stage, tissue rearrangement in the embryos could cause inconsistencies in these three genes. However, because these three genes were newly discovered, evaluating them proved challenging.

### ZENomix limitation

We used ZENomix on *bcd*-knockdown (KD) *D. melanogaster* embryos to test its predictability in mutants with a tissue structure rearrangement. During *Drosophila* development, the anterior-posterior (A-P) axis was created by morphogen gradients of the anterior *bcd* and posterior *nanos* genes. In contrast, in *bcd*-KD embryos, the anterior identity of the embryo was converted to the posterior identity, resulting in a reorganisation of embryonic structural allocation, with loss of the head and thorax^33^ (**Extended Data Fig. 7a**).

First, we attempted to reconstruct *bcd*-KD spatial transcriptomes in a zero-shot manner (**Extended Data Fig. 7b, d**). In this experiment, *bcd*-KD scRNA-seq data from Sakaguchi et al.^34^ and wild-type FISH data (84 genes) from the Berkeley Drosophila Transcription Network Project (BDTNP)^35–37^ were used. We observed that the reconstruction of the *bcd-*KD spatial transcriptomes failed completely. As a control experiment, we used ZENomix to reconstruct *bcd*-KD spatial transcriptomes using the teaching data and *bcd*-KD ISH from Staller et al.^33^ (13 genes) (**Extended Data Fig. 7c, d**). Under these conditions, we confirmed that ZENomix successfully reconstructed *bcd*-KD spatial transcriptomes. To investigate why ZENomix failed under zero-shot conditions, we compared *bcd*-KD spatial transcriptomes with and without teaching data by estimating the original location of the scRNA-seq data point (see **Methods**). In the reconstruction with teaching data (i.e., *bcd*-KD ISH data), the scRNA-seq data points were equally estimated along the A-P axis, whereas most scRNA-seq data points were estimated to be biased toward the posterior region in the reconstruction without teaching data (i.e., wild-type ISH data) (**Extended Data Fig. 7e–g**). These findings indicate that ZENomix incorrectly estimated the origin of the scRNA-seq data points obtained from the anterior region as the posterior region because of anterior and posterior identity conversion.

The A-P axis of *bcd*-KD embryos was folded in half in the gene expression space, whereas the A-P axis of wild-type embryos was clearly distinguished. We reasoned that the A-P conversion of *bcd*-KD embryos could explain the failure of origin estimation (**Extended Data Fig. 7h**). Thus, ZENomix does not reliably predict spatial transcriptomes when the spatial trajectories of wild type and mutant genes in the gene expression space are not correctly calibrated.

## Discussion

We developed ZENomix, a computational framework for reconstructing mutant spatial transcriptomes without teaching data, by introducing the novel zero-shot reconstruction concept. ZENomix recovers spatial information of mutant scRNA-seq data by using the wild-type spatial gene expression atlas as side information and extracting landmark points of spatial coordinates in tissues from the wild-type spatial reference atlas. Using simulated and real scRNA-seq data, we showed that ZENomix reliably predicted the spatial transcriptomes of AD-mutant mouse model and *MZoep*-mutant zebrafish embryos. Furthermore, using this novel, spatially informed screening approach based on ZENomix predictions, we discovered eight novel ND genes in early zebrafish embryos. Therefore, ZENomix is useful for identifying genes with perturbed expression and can provide new insights into the mutant or diseased tissue pathogenesis.

The proposed zero-shot reconstruction method contributes to the field of spatial transcriptomic in two ways. First, it reuses published mutant/disease-specific scRNA-seq data. Reanalysis of public scRNA-seq data can reveal spatial transcriptomes because *in situ* data of mutations or diseases of interest are not required for cross-genotype integration. Since many patient-derived scRNA-seq data have been collected,^38^ zero-shot reconstruction method can be used for wider applications. Second, it is applicable to small, spherical, or dome-like tissues, such as *Drosophila* and zebrafish early embryos, for which ST technology is currently challenging to apply. To the best of our knowledge, only a few studies have revealed the spatial transcriptomes of *Drosophila* and zebrafish embryos using ST technology (Stereo-seq^39^). However, these tissues have been extensively studied using computational reconstruction methods^11,40^. Since the zero-shot method significantly simplifies the computational approach, it can significantly make spatial transcriptomic studies more feasible.

Although many methods using scRNA-seq data have been used to rebuild spatial transcriptomes^11–19^, none can achieve the zero-shot reconstruction of mutant spatial transcriptomes for two reasons. First, the scRNA-seq and *in situ* data are modelled assuming they were obtained under same biological conditions. For example, the neural network-based method gimVI^16^ assumes that *in situ* and scRNA-seq data are generated from a shared latent biological state using a similar function. ZENomix overcomes this bottleneck by modelling mutant and wild type data using different functions based on Gaussian process priors. Second, most previous methods optimised similarity measures (e.g. cosine similarity^18^, mean squared error^19^, inverse correlation^15^, and mutual information^17^) between *in situ* data and reconstructed transcriptomes, making integration of various genotyped data unfeasible. In contrast, ZENomix relies on Bayesian estimation of spatial transcriptomes, enabling reconstruction without any similarity measures.

Our newly developed screening approach, based on ZENomix prediction, is critical to identify genes whose expression is perturbed in mutant tissues. Our screening approach can use data related to gene expression changes in specific regions between the wild type and the mutant (e.g. the embryonic margin in *MZoep*), in contrast to bulk transcriptomics-based screening using microarray and RNA-seq. Using our spatially informed screening method, we discovered previously unknown eight new Nodal-downregulated genes. We also identified 74 putative NU genes, which was consistent with the previous bulk microarray-based screening^30^. These findings highlight the significance of spatial information in knockdown/knockout analyses, suggesting that ZENomix can identify new biological mechanisms through zero-shot reconstruction.

This study had some limitations. Although ZENomix could be applied for the two mutant tissues, we cannot conclude it can be applied to any mutant tissue for two reasons. First, this is based on the assumption that the morphology of the mutant tissue des not differ significantly from that of the wild type. Second, only gene expression of the scRNA-seq data points was used to assign spatial locations. ZENomix failed to reconstruct the spatial transcriptomes of *bcd*-KD *Drosophila* embryos, which have conversion of A-P axis from wildtype embryos. Therefore, studies on modelling the differences in tissue morphology between wild types and mutants, focusing on the changes in tissue morphology, can address this challenge. In addition, incorporating prior information into the gene regulatory network may refine the prediction of the original location of the mutant scRNA-seq data.

## Methods

### Zebrafish

The wild-type strain RIKEN WT (RW) was used in this study. All zebrafish experiments were approved by the Animal Studies Committee of the Nara Institute of Science and Technology.

### Whole-mount ISH

Whole-mount ISH was conducted as described previously^41,42^. Briefly, PCR was used to generate template DNAs for *axin2, cdx4, cxcr4a, eve1, foxd5, gbx1, msx1b, sox19a, sp5l*, and *szl* using forward and reverse primers with T3 and T7 sequences, respectively, inserted at the 5’ ends; the T3 and T7 primers were used to confirm the sequence. pCRII-*cxcr4b* was used as the template, and T7 or SP6 RNA polymerases were used to synthesise DIG antisense RNA probes.

### lefty1 overexpression in zebrafish embryo

*lefty1* mRNAs were synthesised using the mMessage mMachine SP6 transcription kit (Thermo Fisher Scientific) with pCS2-*lefty1* (kindly gifted by Dr. Masashiro Hibi) as the template^32^. Then, 5 pg of the synthesised *lefty1* mRNAs was injected into one-cell stage zebrafish embryos, and the embryos were used for ISH.

### ZENomix model

ZENomix uses wild-type spatial reference data and mutant scRNA-seq data as inputs. In spatial reference data (e.g. ST data) generated using *in situ* methods, gene expression vectors are available for all regions/cells whose locations in the tissue are known. The gene expression vector of gene *j* is represented as 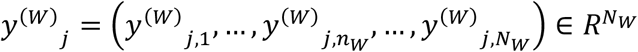, where cells are indexed by *n*_*W*_(*n*_*W*_ ∈ {1, 2,…, *N*_*W*_}), and *N*_*W*_ is the total number of cells in the tissue of interest.

By contrast, in the scRNA-seq dataset of the genotype of interest, gene expression vectors lack information regarding location of cells in the tissue. The expression vector of gene *j* is represented as 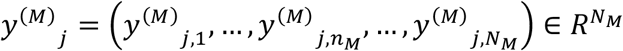, where cells are indexed by *n*_*M*_ (*n*_*M*_ ∈ {1, 2, …, *N*_*M*_}), and *N*_*M*_ is the total number of cells used for scRNA-seq measurement. The number of dimensions of these two data points *p* is the same because we considered landmark genes.

#### Generative model

The scRNA-seq and spatial reference data were modelled as a generative model.

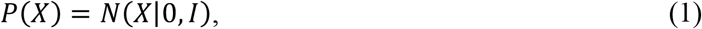

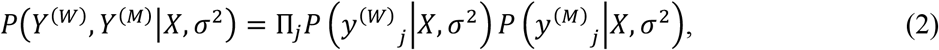

where *N*(0, *I*) represents a standard Gaussian distribution with *q* dimensions; 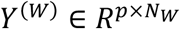 and 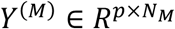 indicate *in situ* and scRNA-seq data matrices, respectively; *X* indicates the common latent variable matrix (spatial landmark points) with dimension *q*; and *h* indicates the genotype or modality of data (*h* ∈ {W (wild-type spatial reference data), M (mutant scRNA-seq data)}). The following functions of low-dimensional latent variables were used to generate gene expression:

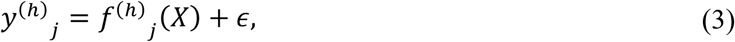

where *ϵ* indicates the Gaussian observation noise with zero mean and standard deviation *σ*. Note that *σ* is common among the data modalities. The aim was to estimate the posterior distribution of the latent variables, denoted as *P*(*X*|*Y*^(*W*)^, *Y*^(*M*)^).

To assume a nonlinear transformation from latent variables to gene expression levels, the Gaussian process prior was placed on the projection function *f*^(*h*)^_*j*_ as follows:

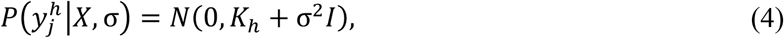

where *K*_*h*_ is the gram matrix defined by the kernel function *k*(*x*^(*h*)^, *x*′^(*h*)^; θ_*k*_) between two distinct latent variables, *x*^(*h*)^ and *x*′^(*h*)^ with kernel hyperparameters, *θ*_*k*_, and *x*^(*h*)^ is a latent variable corresponding to each observation, *y*^(*h*)^.

#### Inference scheme

The ZENomix model was established to estimate the posterior probabilities of shared latent variables (spatial landmark points) *P*(*X*|*Y*^(*W*)^, *Y*^(*M*)^). However, because the two datasets are unpaired, it was difficult to estimate the latent variables shared by *Y*^(*W*)^ and *Y*^(*M*)^, implying that we cannot use the methods for paired data (e.g. canonical correlation analysis^43^ or multimodal mixture-of-expert variational autoencoders^44^). To solve this, vGPLVM-MMD, a new inference scheme that extracts spatial landmark points from wild-type data, was proposed.

The vGPLVM-MMD scheme comprises two parts; the first part is to independently map *Y*^(*W*)^ and *Y*^(*M*)^ onto a low-dimensional latent space, and the second part is to match the latent data distributions. In the first part, the generative model is separated into two distinct models: *Y*^(*W*)^ and *Y*^(*M*)^, where *X*^(*h*)^ is a separate latent variable matrix corresponding to each data matrix *Y*^(*h*)^. Consequently, gene expression is generated by the function of the low-dimensional latent variable *X*^(*h*)^, which varies for each data point, as shown below:

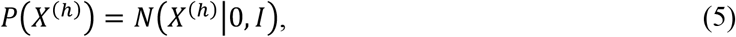

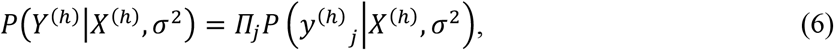

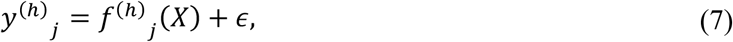

where the Gaussian process prior is placed on the projection function *f*^(*h*)^_*j*_. From this model, we can estimate the posterior distributions *P*(*X*^(*h*)^|*Y*^(*h*)^) for each data. In practice, we used variational inference to estimate the approximated posterior distribution *q*(*X*^(*h*)^), rather than *P*(*X*^(*h*)^|*Y*^(*h*)^). *q*(*X*^(*W*)^) and *q*(*X*^(*M*)^) exhibit different distributions in this model.

This model (Eq. 5–7) is known as the Bayesian GPLVM^24^. As the posterior distribution is analytically intractable in this model, Titsias and Lawrence^24^ and Damianou et al.^25^ developed a variational GPLVM using inducing inputs. Following their methods, the generative model can be described as follows:

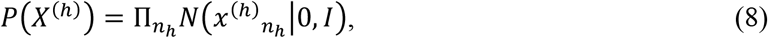

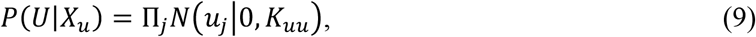

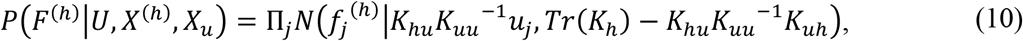

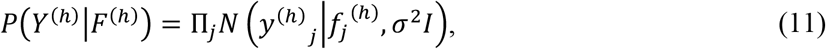

where 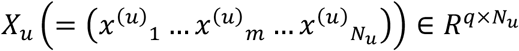 and 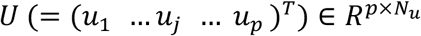 indicate the induced inputs in the latent space and extra samples in the observation space, respectively, and *F*^(*h*)^ indicates the GP mapping values. In Eq. (10), the gram matrices 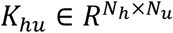 and 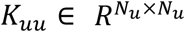 are defined as follows:

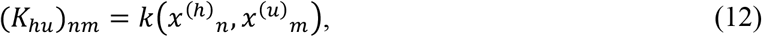

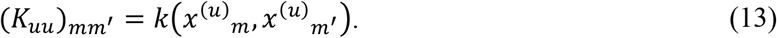

Using a Gaussian distribution, the variational inference is used to approximate the true posterior *P*(*X*^(*h*)^|*Y*^(*h*)^) as follows:

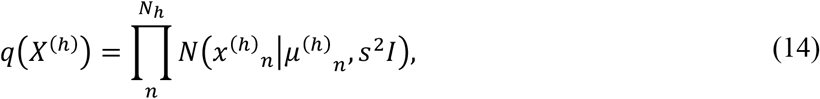

where s indicates the noise intensity, which is shared by all distributions. The lower bound of *P*(*Y*^(*h*)^)

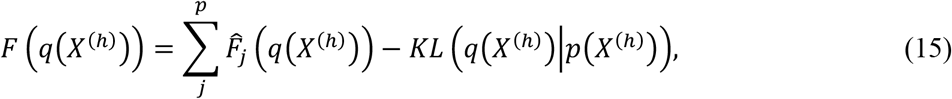

is expressed as follows:

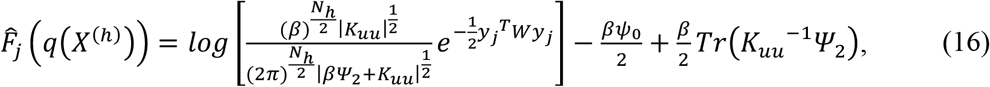

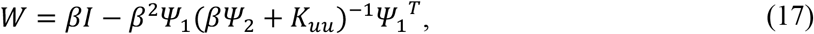

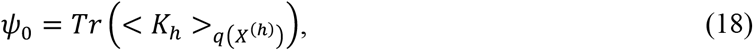

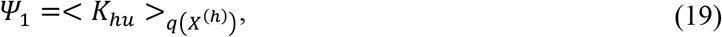

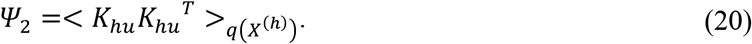

where <·>_*p*(*x*)_ denotes expectation under the distribution *p*(*x*), and *KL*(*q*(*x*) || *p*(*x*)) indicates the Kullback–Leibler (KL) divergence between distributions *q*(*x*) and *p*(*x*).

In our implementation, *Ψ* statistics (*ψ*_0_, *Ψ*_1_, and *Ψ*_2_) were computed using the Gaussian–Hermite approximation, as in Gpy^45^. The latent representation of each data point can be obtained by maximising the variational bound for each genotype, h.

In the second part of the vGPLVM-MMD scheme, the independently derived posterior distributions *q*(*X*^(*W*)^) and *q*(*X*^(*M*)^) are combined, such that the two distributions may be nearly matched as *q*(*X*^(*W*)^) ≈ *q*(*X*^(*M*)^) to extract the common latent variable *X*. To make the two distributions almost identical, the distance between the two distributions, *q*(*X*^(*W*)^) and *q*(*X*^(*M*)^), was minimised. MMD^23^ was used to compute the distance (see the subsequent *MMD calculations* section).

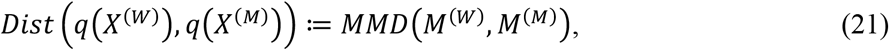

where 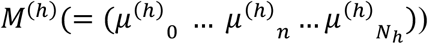 is the mean of *q*(*X*^(*h*)^).

#### MMD calculations

MMD is the nonparametric distance between two sample distributions embedded in a Reproducing Kernel Hilbert Space (RKHS). Let us assume a general situation where two data samples, *X* and *Y*, have identical dimensions. The empirical estimate of the MMD between the data points *X* and *Y* is as follows:

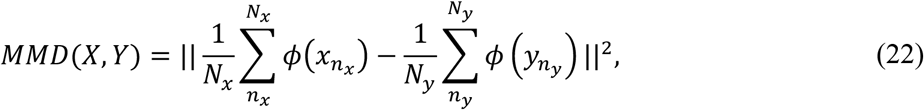

where *ϕ* indicates the kernel-induced feature map. By expanding Eq. (22) and replacing the inner products with their kernel values (the kernel trick), MMD is given as

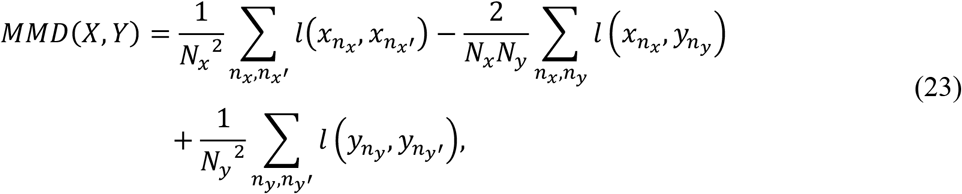

where l(x, y) denotes the kernel function with MMD hyperparameters, *θ*_*l*_. When the RKHS is universal, the MMD approaches zero asymptotically if and only if the two distributions are the same^23^.

#### Variational inference

To estimate the parameters and hyperparameters of ZENomix, we integrated the first and second parts of the vGPLVM-MMD. Thus, the cost function of the vGPLVM-MMD can be denoted as follows:

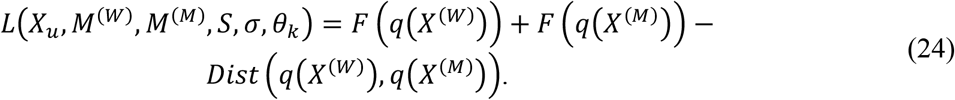

This cost function was maximised using a gradient-based optimisation algorithm (see the *ZENomix implementation* section) to extract the common latent variables of the two datasets. The MMD hyperparameters *θ*_*l*_ are fixed in ZENomix. The computational cost of evaluating *F* (*q*(*X*^(*h*)^)) is *O*(*N*_*h*_ ∗ *N*_*u*_^2^) and that of *MMD*(*M*^(*W*)^, *M*^(*M*)^) is *O*((*N*_1_ + *N*_*R*_)^2^). If 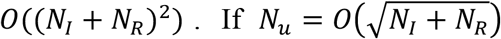, the computational complexity of evaluating L is *O*((*N*_1_ + *N*_*R*_)^2^).

#### Spatial reconstruction

In the second step of ZENomix, the mutant spatial gene expression profiles of gene *j* are obtained by mapping the latent variables of wild-type *in situ* data, *X*^(*W*)^, to mutant scRNA-seq space as *F*^∗(*M*)^_*j*_ ≔ *f*^(*M*)^_*j*_ (*X*^(*W*)^). The posterior distribution of *F*^∗(*M*)^_*j*_ can be inferred using Eq. (8–11) as follows:

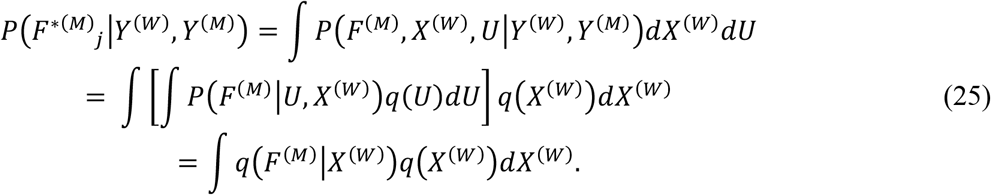

Although this integral is analytically intractable, Titsias and Lawrence and Damianou et al. showed that the mean and covariance of *F*^∗(*M*)^_*j*_ can be calculated as follows:

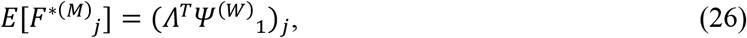

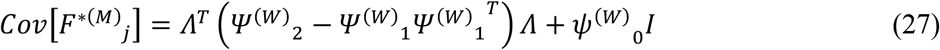

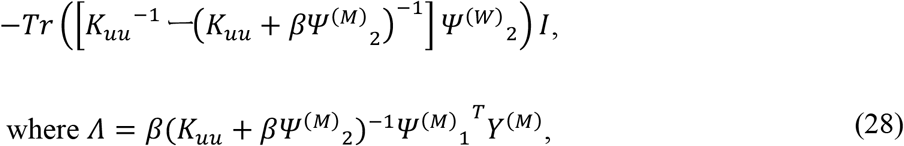

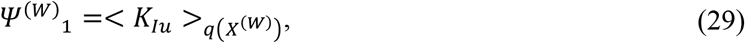

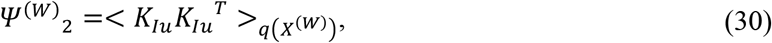

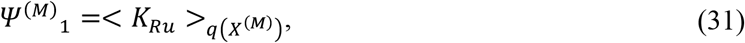

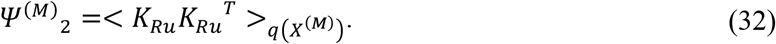

#### ZENomix implementation

In our implementation, we selected the Matern 3/2 kernel for both kernel functions *k*(*x, y*) and *l*(*x, y*) with length scales of 1/*σ*^*f*^_*k*_(= *θ*_*k*_) and 1/*σ*^*f*^_*l*_(= *θ*_*l*_), respectively. To initialise the mean of the posterior distributions *M*_*l*_ and *M*_*R*_, principal component analysis (PCA) was used for the input data. Before the first step of ZENomix, all input data were z-scored, and ∑_*h*_ *F* (*q*(*X*^(*h*)^)) and *MMD*(*M*^(*W*)^, *M*^(*M*)^) were divided by their initial values for normalisation. **Supplementary Table 2** shows the detailed initial parameter settings. The L-BFGS-B algorithm implemented in SciPy was used as the gradient-based optimisation method.

### Data collection and pre-processing

#### Mouse OB data

Wild-type OB ST data were originally generated by Ståhl et al.^21^. Normalised data and a list of highly variable genes were downloaded from SpatialDB^46^ (http://www.spatialomics.org/SpatialDB/), and Rep11 was used for further analyses. The mouse AD OB ST dataset was downloaded from Mendeley Data^3^ (https://doi.org/10.17632/6s959w2zyr.1). Normalised mouse wild-type and AD data were used for further analyses (including pre-processing and integration with wild-type ST data). To integrate the wild-type ST data with the AD ST data, the spatial information of the AD ST data was ignored, and each spot was considered a simulated scRNA-seq data point. We selected highly variable genes from the wild-type reference downloaded from SpatialDB as landmark genes. Genes not included in the simulated scRNA-seq data or whose expression in the simulated scRNA-seq data was 0 were removed from the landmark genes measured in ISH data.

#### Zebrafish early embryo data

Zebrafish early embryo ISH data were downloaded from the Satija Lab homepage (https://satijalab.org/). For zebrafish early embryo scRNA-seq data, we downloaded the raw data from the Gene Expression Omnibus database (accession number GSE106587) and pre-processed them as previously described^9^ for both genotypes (wild type and *MZoep*). Genes not included in the scRNA-seq data or whose expression in the scRNA-seq data was 0 were removed from the landmark genes measured in ISH data. The geometric data were obtained from Cang et al.^47^.

#### Drosophila melanogaster embryo data

For the wild-type reference data, we used the modified ISH data generated by Sakaguchi et al.,^34^ originally obtained from BDTNP (D_mel_wt atlas_r2.vpc from http://bdtnp.lbl.gov) and DVEX (bdtnp.txt). For the *bcd*-KD reference data, we downloaded *bcd*-KD ISH data^33^ from FigShare (https://figshare.com/articles/dataset/A_gene_expression_atlas_of_a_bicoid_depleted_Drosophila_e_mbryo/1270915) and used the cohort name 5:76–100 (the end of stage 5) as the reference. The ISH data were log-scaled before the ZENomix procedure. Wild-type and *bcd*-KD scRNA-seq data were obtained from Sakaguchi et al.^34^. Both scRNA-seq datasets were pre-processed as previously described^34^.

### ZENomix and downstream analysis

ZENomix was built in Python 3.10.5 and is available on GitHub (https://github.com/yasokochi/ZENomix). All other software used in the ZENomix are publicly available: Numpy version 1.22.4 (https://numpy.org/) for calculation; scipy==1.9.3 (https://scipy.org/) for calculation; jax version 0.3.25 (https://jax.readthedocs.io/en/latest/index.html) for calculation; scikit–learn version 1.1.3 (https://scikit-learn.org/) for traditional machine learning (e.g. PCA); and pandas version 1.5.1 (https://pandas.pydata.org/) for reading data frames. For downstream analysis, we used Scanpy version 1.9.1^48^ (https://scanpy.readthedocs.io/en/stable/) for scRNA-seq data analysis and Squidpy version 1.2.3^49^ (https://squidpy.readthedocs.io/en/stable/) for spatial transcriptome data analysis.

### Calculating Moran’s I statistics

The accuracy of the predicted spatial transcriptomes was evaluated by comparing spatial autocorrelations (Moran’s I values) between the predicted and original spatial transcriptome data. Moran’s I value was calculated as follows:

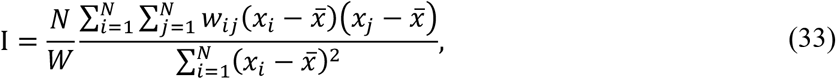

where *N* is the number of data points indexed by *i* and *j*; *x* depicts the data point; 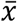 is the mean of *x*; *w*_*ij*_ is a matrix of spatial weights with zeroes on the diagonal; and *W* is the sum of all *w*_*ij*_. We used the *gr*.*spatial_autocorr* function with mode = ‘moran’ in Squidpy for implementation.

### Selecting spatially DE genes

We first calculated the maximum absolute value and standard deviation of expression differences between *MZoep*-mutant spatial transcriptomes and those of the wild type. Subsequently, 142 genes were selected by manual thresholding of 8 > *max_diff* > 2 and *std* > 0.72. Based on the mean expression changes in the embryo margin, these genes were classified into two groups: putative NU and ND genes. The embryo margin was defined as tier = ‘1–2’ or ‘3–4’ in the zebrafish geometry data. The spatial gene expression profiles of the putative NU and ND genes were clustered via hierarchical clustering using the *cluster*.*hierarchy*.*fcluster* function in SciPy based on the correlation matrices of the gene expression changes, and four and three modules were obtained for the putative NU and ND genes, respectively. As module 1 of the putative NU genes and modules 1 and 3 of the putative ND genes showed a universal gene expression change, 87 spatially DE genes (74 and 13, respectively) were excluded.

### Data visualisation

The publicly available tools Matplotlib version 3.6.2 (https://matplotlib.org/) and Seaborn version 0.12.1 (https://seaborn.pydata.org/) were used to visualise the data. For mouse OB data, we used the spatial location data of the ST data of the collected wild-type and AD-mutant mouse OB data as described above in the *Data collection and pre-processing* section. To visualise the zebrafish embryo data, we used the plot_zf function created by Cang et al.,^47^ which interpolates the original 64 data points *in situ*. For the *Drosophila* embryo data, the embryos were visualised as previously described^17^.

## Data Availability

This study is a reanalysis of existing data. The websites from which the data were collected are mentioned in the ***Data collection and pre-processing*** subsection of the **Methods** section.

## Code Availability

ZENomix is developed under Python 3.10.5 and is available on GitHub (https://github.com/yasokochi/ZENomix).

## Acknowledgements

We are grateful to Prof. Masahiko Hibi and Dr. Ken Nakae for their valuable discussions, Ms. Maiko Yokouchi for technical assistance, and Prof. Masahiko Hibi for kindly gifting pCS2-*lefty1*. We are also thankful for the research opportunities provided by Prof. Michiyuki Matsuda and the Medical Scientist Training Program of Kyoto University (to Y.O.). This study was supported in part by the Moonshot R&D–MILLENNIA Program (grant number JPMJMS2024-9 to H.N.) of the Japan Science and Technology Agency (JST), Grant-in-Aid for Transformative Research Areas (B) (grant number 21H05170 to H.N.), Grant-in-Aid for Scientific Research (B) (grant number 21H03541 to H.N.) from the Japan Society for the Promotion of Science (JSPS), and Cooperative Study Program of Exploratory Research Center on Life and Living Systems (ExCELLS; program number 19-102 to H.N.), and Grant-in-Aids for Scientific Research (B) and Challenging Exploratory Research (grant numbers 22H02821 and 21K19265 to T.M.) from the JSPS.

## Author Contributions

Y.O. and H.N. conceived the project. Y.O. developed the model. Y.O. and T.M. conducted the experiments. Y.O., S.S., and T.K. analysed the data. Y.O. and H.N. wrote the manuscript with input from all the authors.

## Supplementary Information

Supplementary information is available for this paper.

## Extended Data Figures

**Extended Data Fig. 1.**
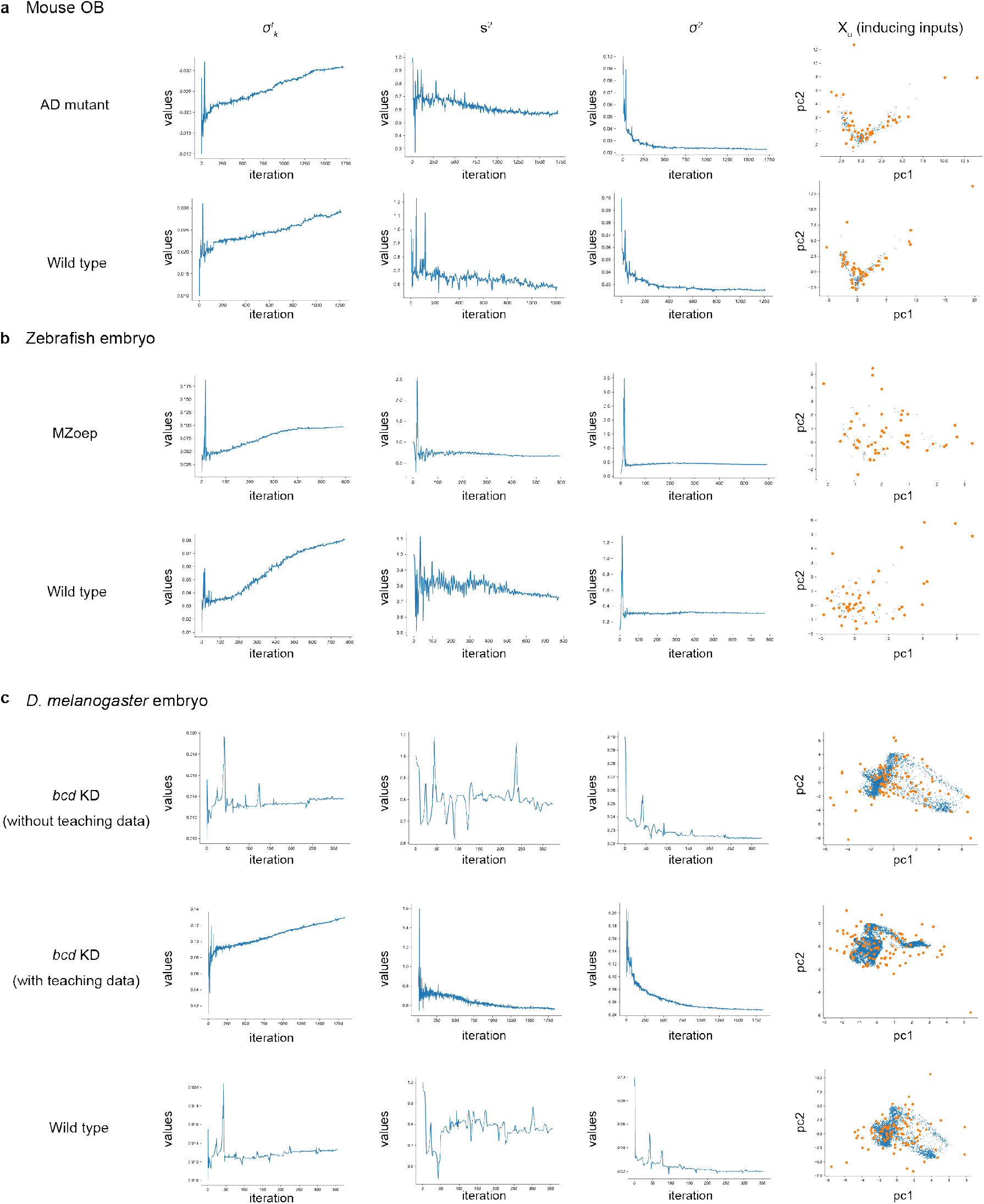
ZENomix parameter convergence. Parameter values during the first step of ZENomix for the prediction of (**a**) mouse OB, (**b**) zebrafish embryo, and (**c**) *D. melanogaster* embryo spatial transcriptomes. Each parameter is described in the **Methods** section. Principal component analysis was used to show the final placements of the inducing points. The large orange and small blue points indicate the induced and spatial landmark points, respectively.

**Extended Data Fig. 2.**
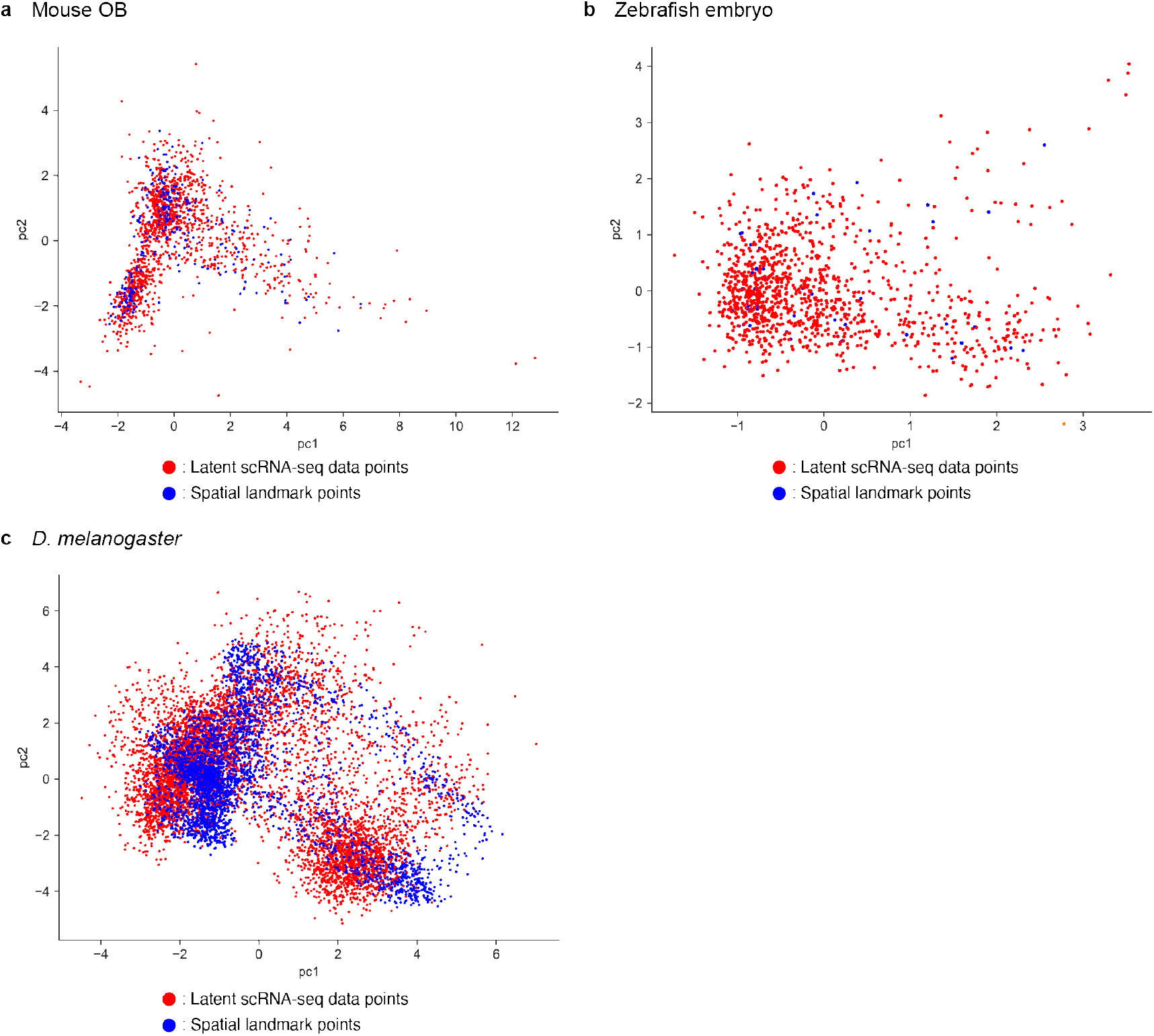
Latent data calibration for wild-type prediction. Scatter plots of calibrated distributions of spatial landmark and scRNA-seq data points (**Fig. 2b**) for prediction of (**a**) wild-type mouse OB, (**b**) wild-type zebrafish embryo, and (**c**) wild-type *D. melanogaster* embryo spatial transcriptomes. Principal component analysis was used to visualise the latent space.

**Extended Data Fig. 3.**
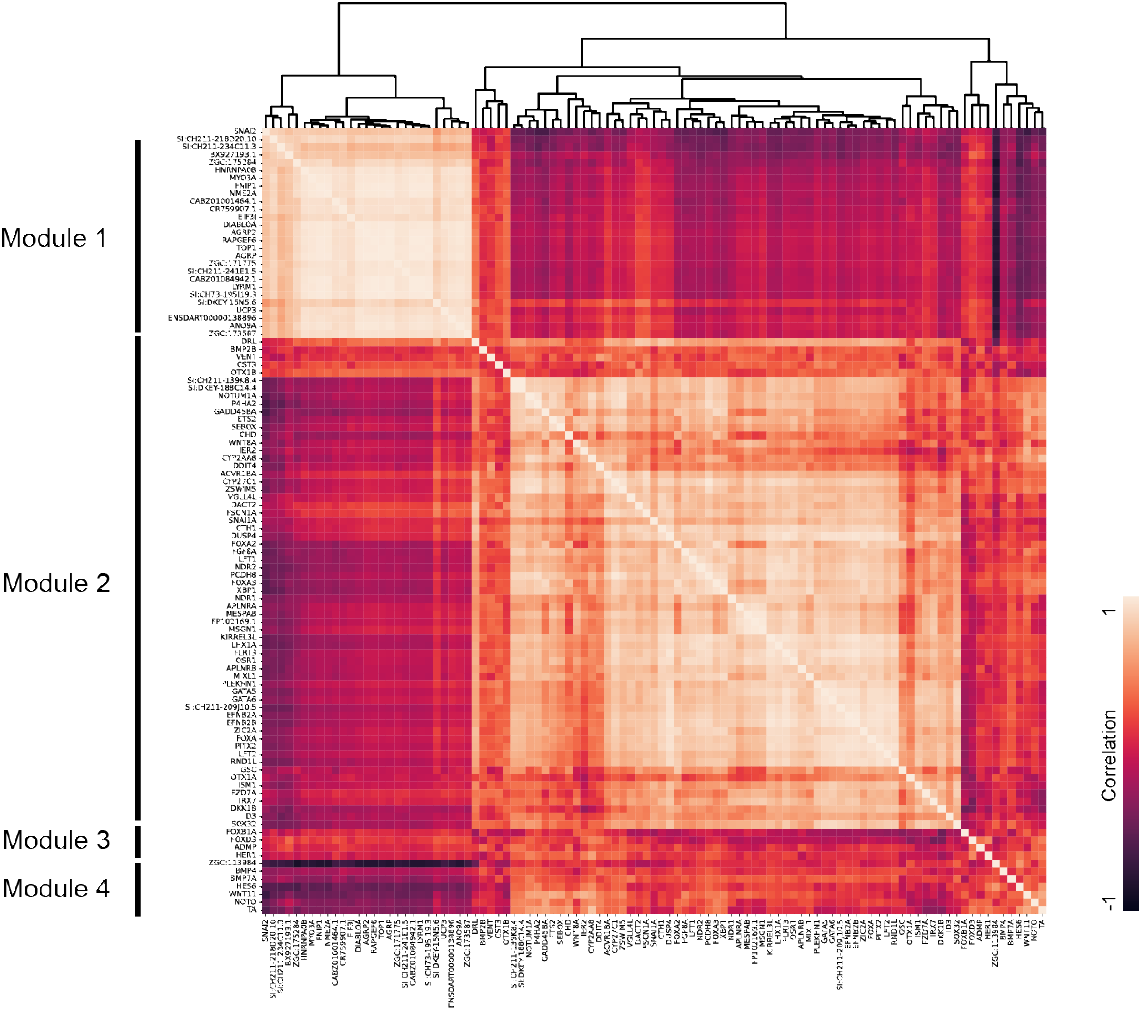
Putative NU gene hierarchical clustering. (a) Hierarchical clustering of the putative NU genes (corresponding to **Fig. 5d**). The heatmap indicates the correlations among the changes in the expression of newly screened NU genes.

**Extended Data Fig. 4.**
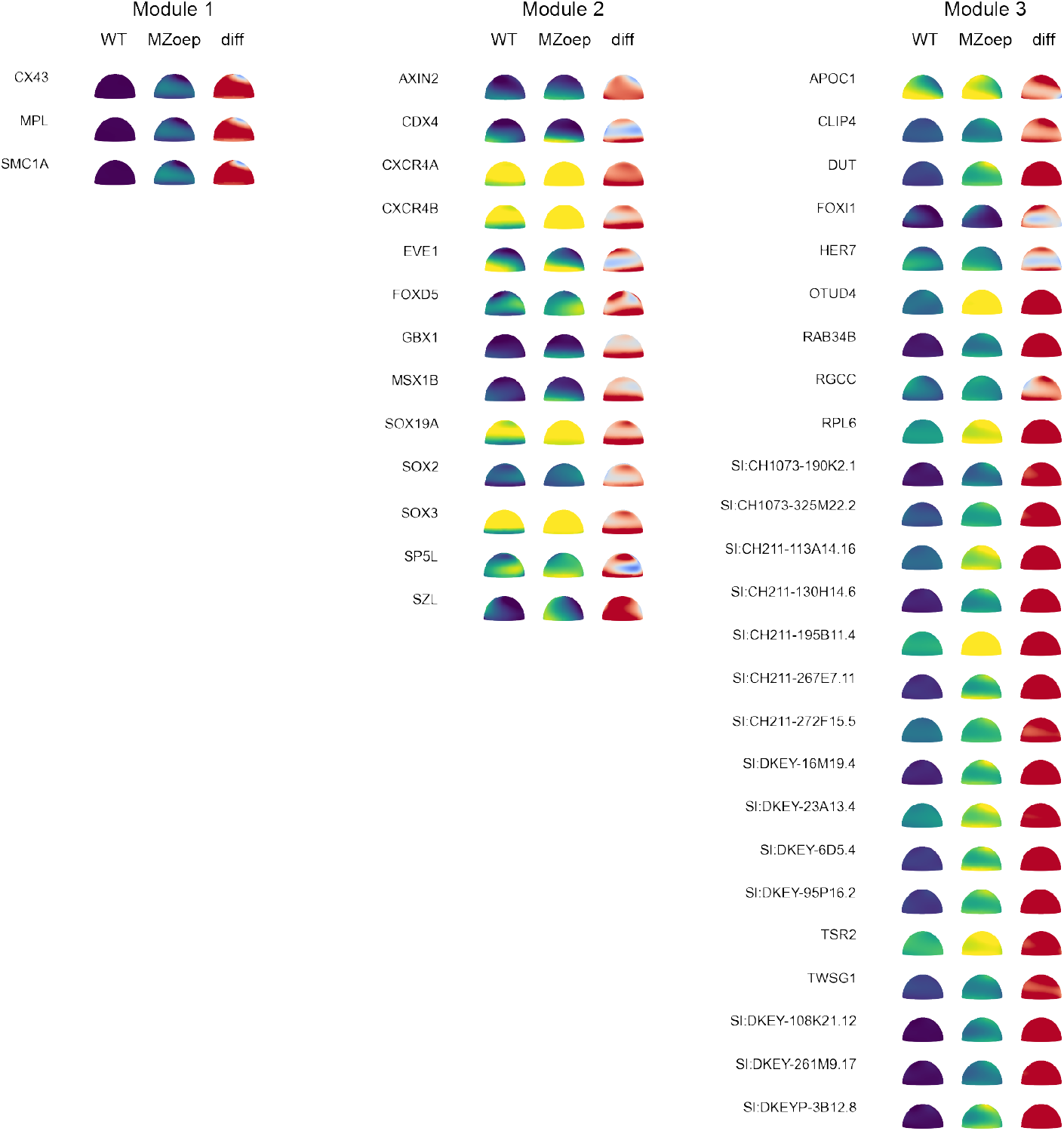
All putative NU genes. The predicted spatial gene expression patterns of wild-type and MZoep zebrafish embryos for all putative NU genes. ‘diff’ indicates the expression difference between the MZoep and wild-type spatial transcriptomes.

**Extended Data Fig. 5.**
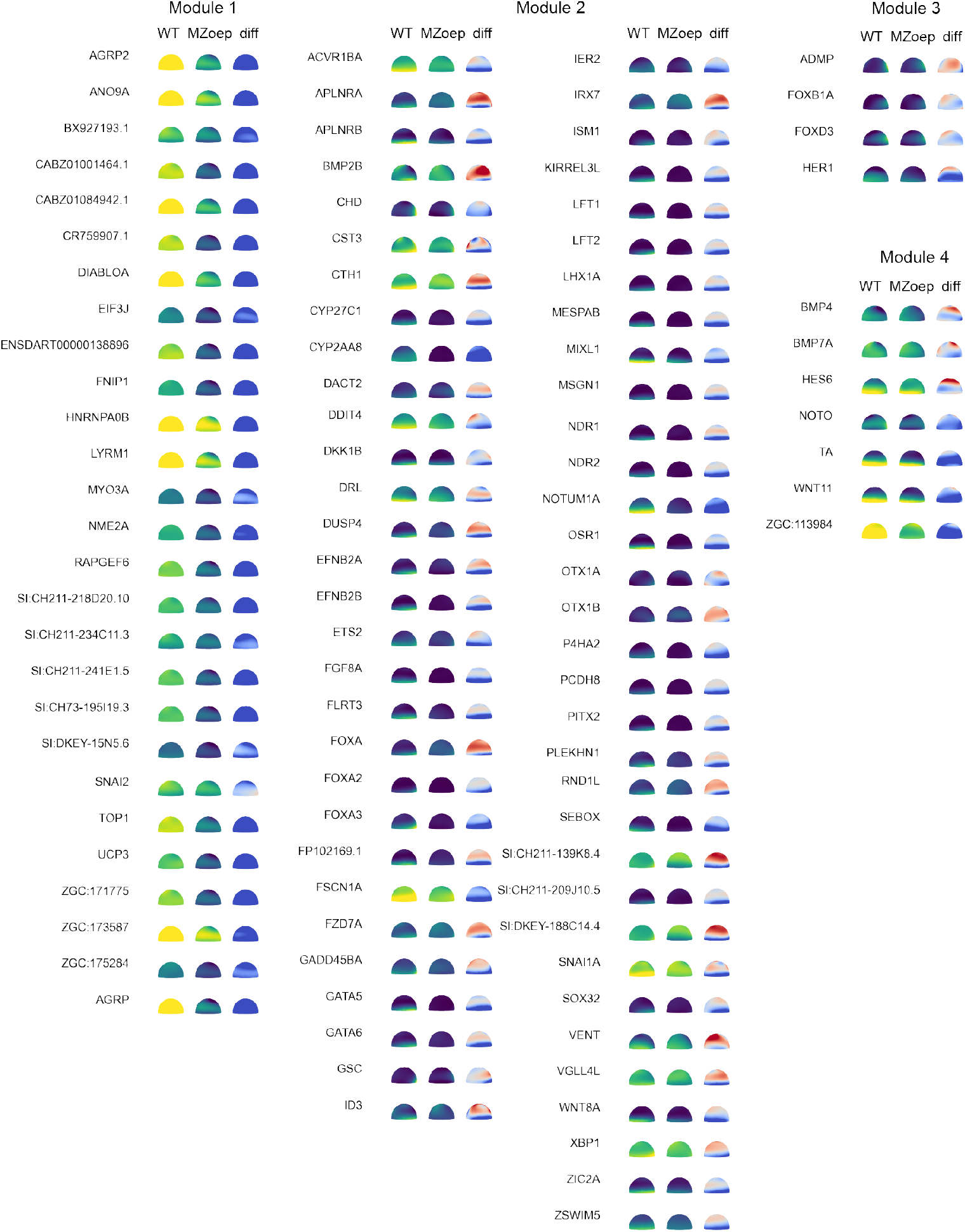
All putative ND genes. Predicted spatial gene expression patterns in wild-type and MZoep zebrafish embryos for all putative ND genes. ‘diff’ indicates the expression difference between the MZoep and wild-type spatial transcriptomes.

**Extended Data Fig. 6.**
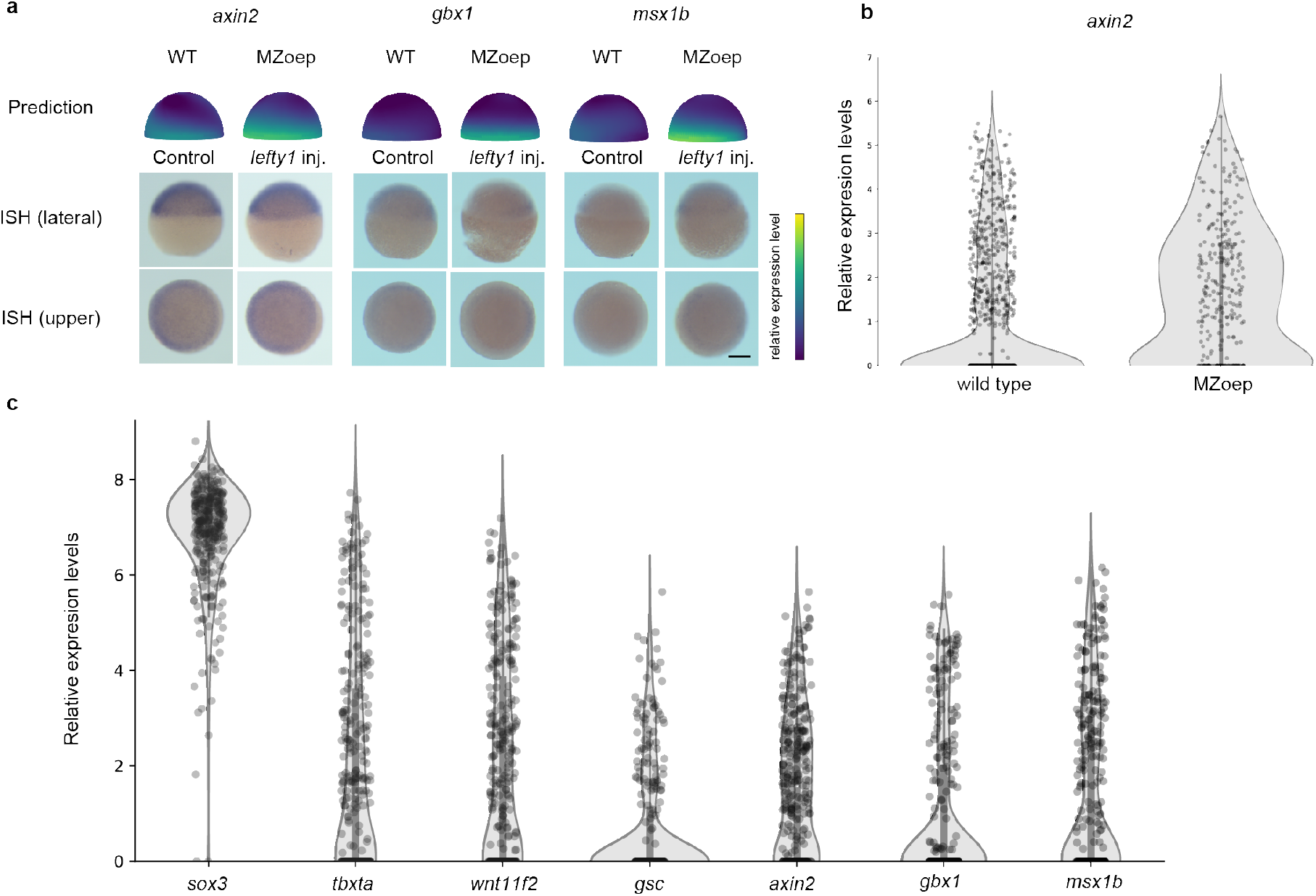
Putative ND genes inconsistent with those of ISH experiments. **a**, Whole-mount ISH experiments for *axin2, gbx1*, and *msx1b*. From top to bottom: ZENomix-predicted expression pattern, ISH lateral view, and upper view of ISH. For each gene, wild-type and mutant expression patterns are displayed (ZENomix prediction: wild-type and *MZoep*-mutant embryos; ISH experiment: control and *lefty1*-injected embryos). Scale bar, 200 μm. **b**, Violin plot showing the differences in *axin2* expression between wild type and MZoep scRNA-seq data. scRNA-seq data showed that MZoep embryos had higher *axin2-*expression levels than did wild-type embryos. **c** Violin plots of *axin2, gbx1*, and *msx1b* expression levels. *sox3, tbxta, wnt11f2*, and *gsc* expression levels are shown in the references. *axin2, gbx1*, and *msx1b* are moderately expressed when compared to genes showing low expression (*gsc*) and those showing high expression (*sox3, tbxta*, and *wnt11f2*). Fig. 4 shows the ISH images of *gsc, sox3, tbxta*, and *wnt11f2*.

**Extended Data Fig. 7.**
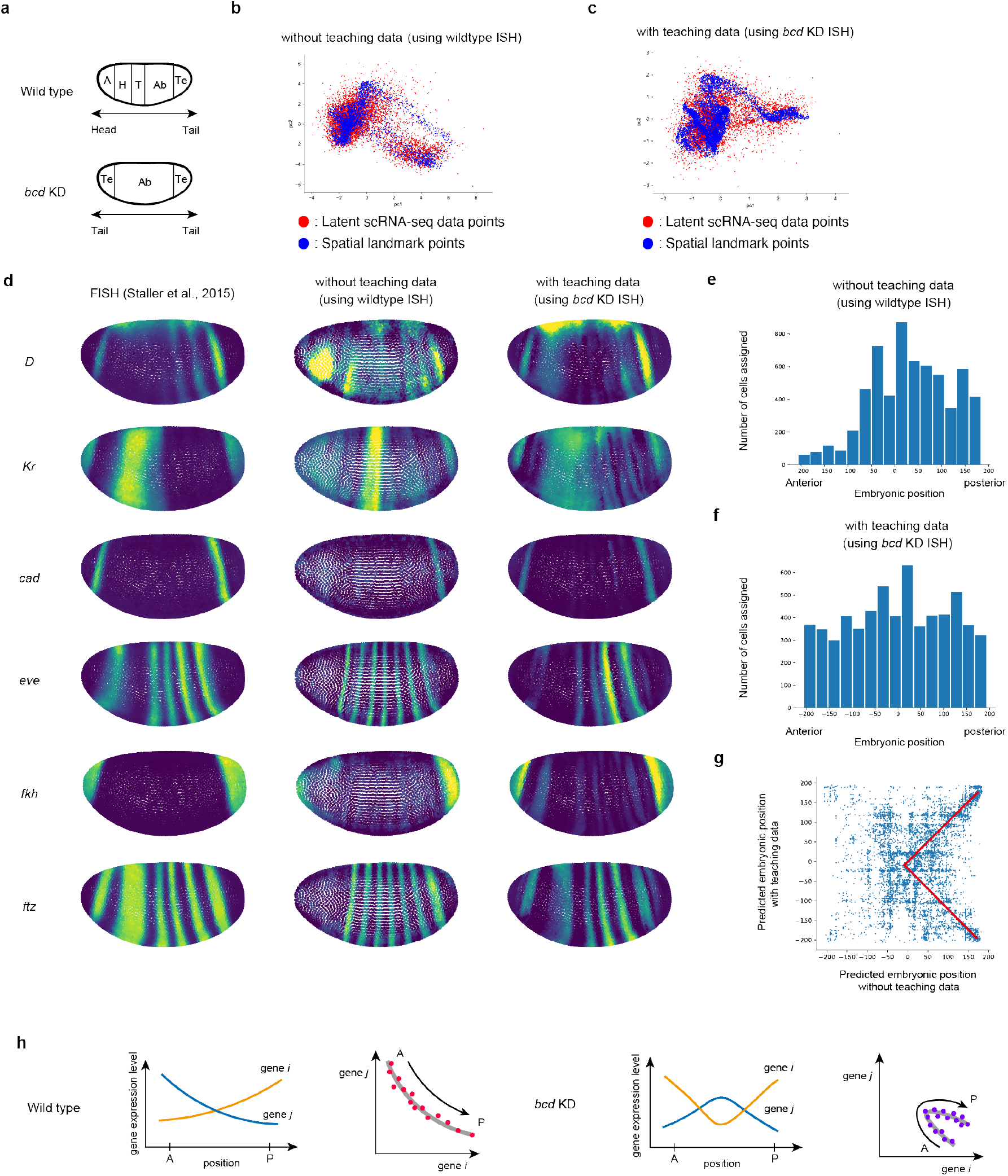
ZENomix application to *bcd*-KD *Drosophila* embryos. **a**, Wild-type and *bcd*-KD *Drosophila* embryo phenotypes. The anterior region of the KD embryo is lost and converted to the posterior region. ‘A’, ‘H’, ‘T’, ‘Ab’, and ‘Te’ indicate the acron, head, thorax, abdomen, and telson, respectively. **b, c** Scatter plots of matched data point distributions of (**b**) wild-type spatial reference and *bcd*-KD scRNA-seq data and (**c**) *bcd*-KD spatial reference and *bcd*-KD scRNA-seq data (corresponding to **Fig. 2b**). Principal component analysis was used to visualise shared latent spaces. (**d**) The *bcd*-KD embryo spatial transcriptome experiment and prediction. We used the FISH data of *bcd*-KD embryos reported by Staller et al. for the experimental data^33^. **e, f** Estimated origin of *bcd*-KD scRNA-seq data points when referencing (**e**) wild type and (**f**) *bcd*-KD *in situ* data. The x- and y-axes indicate the estimated embryonic position and number of data points, respectively. (**g**) Relationship between the estimated origin of *bcd*-KD scRNA-seq data points when referencing the *in situ* data of wild-type (**e**) and *bcd*-KD embryos (**f**). The red line structure indicates the symmetric conversion of embryonic structures in the *bcd*-KD embryo, as depicted in **a. h**, The Simple, one-dimensional model of wild type and *bcd*-KD tissues. The blue and orange lines indicate genes i and j expression profiles, respectively. In the gene expression space, the mutant trajectory corresponds to the posterior part of the wild-type trajectory.

## Supplementary Tables

**Supplementary Table 1.**
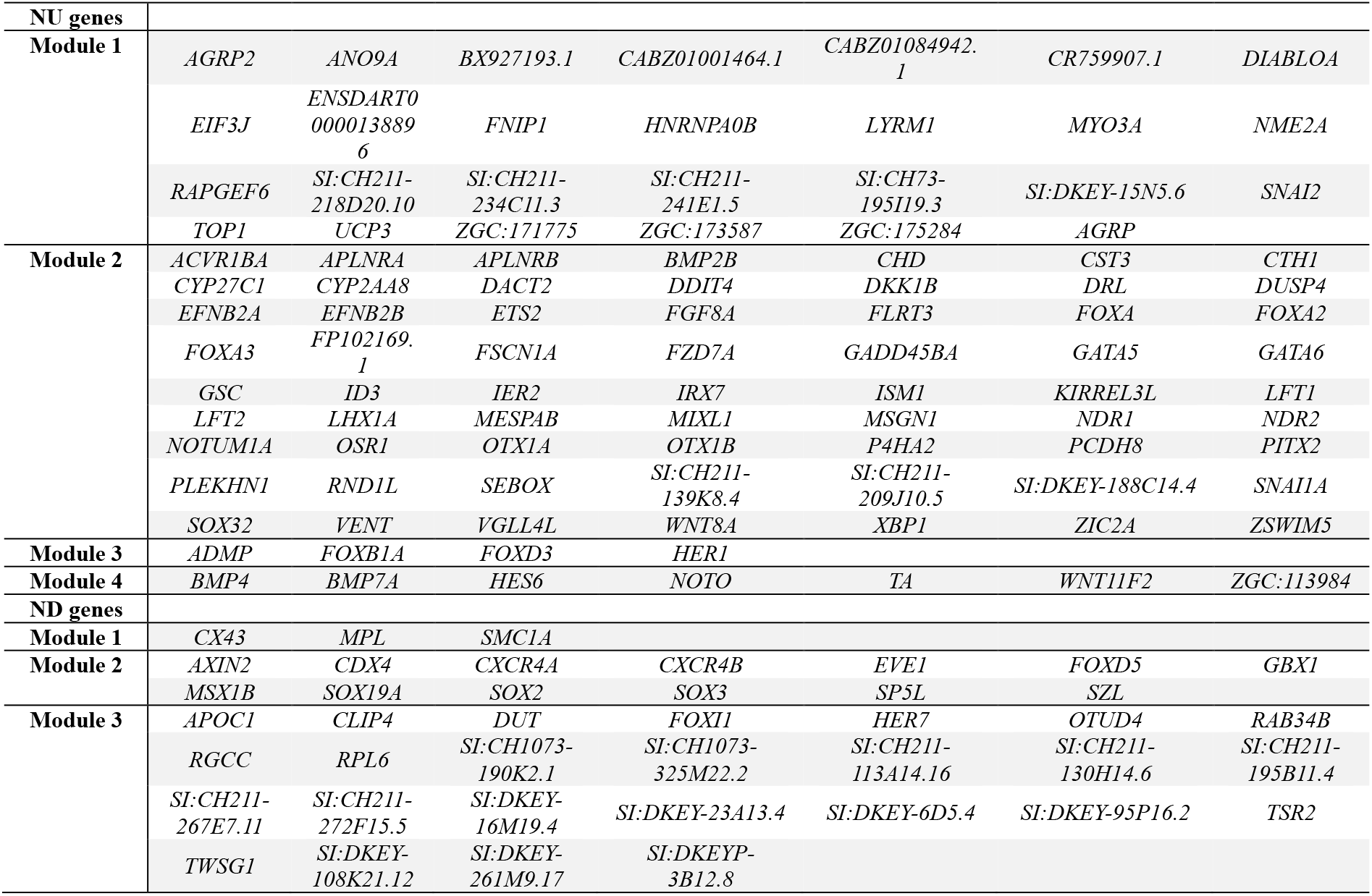
Putative NU and ND genes.

**Supplementary Table 2.**
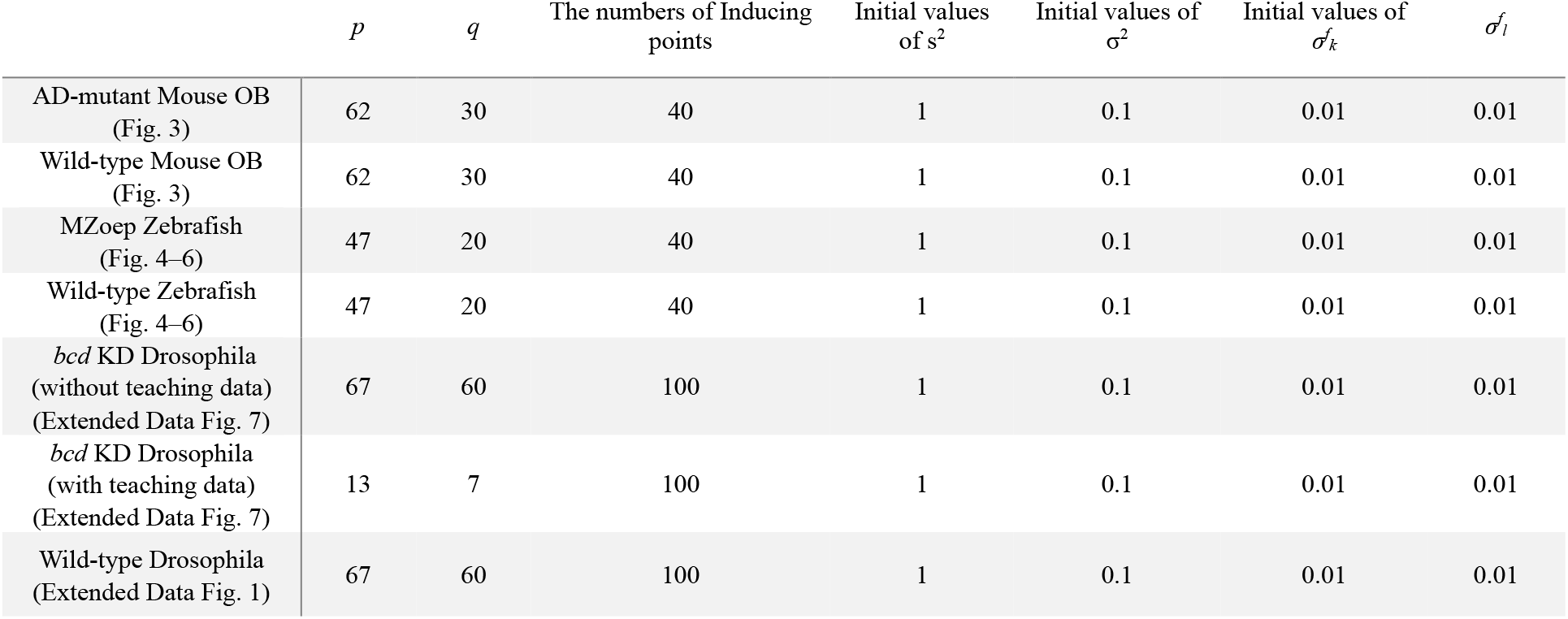
Parameter values used in this study.

